# Multiscale modelling shows how cell-ECM interactions impact ECM fibre alignment and cell detachment

**DOI:** 10.1101/2024.12.05.627121

**Authors:** Juan Arellano-Tintó, Daria Stepanova, Helen M. Byrne, Philip K. Maini, Tomás Alarcón

**Affiliations:** Centre de Recerca Matemàtica, Bellaterra (Barcelona), Spain; Departament de Matemàtiques, Universitat Autònoma de Barcelona, Bellaterra (Barcelona), Spain; Wolfson Centre for Mathematical Biology, Mathematical Institute, University of Oxford, Oxford, UK; Ludwig Institute for Cancer Research, Nuffield Department of Medicine, University of Oxford, Oxford, UK; Institució Catalana de Recerca i Estudis Avançats (ICREA), Barcelona, Spain; Barcelona Collaboratorium for Theoretical & Predictive Biology, Barcelona, Spain

## Abstract

The extracellular matrix (ECM) is a dynamic network structure that surrounds, supports, and influences cell behaviour. It facilitates cell communication and plays an important role in cell functions such as growth and migration. One way that cells interact with the ECM is via focal adhesions, which enable them to sense and respond to matrix mechanical properties and exert traction forces that deform it. This mechanical interplay between cells and the ECM, many aspects of which remain incompletely understood, involves the coordination of processes acting at different spatial scales and is highly influenced by the mechanical properties of the cells, ECM and focal adhesion components. To gain a better understanding of these mechanical interactions, we have developed a multiscale agent-based model based on a mechanical description of forces that simultaneously integrates the mechanosensitive regulation of focal adhesions, cytoskeleton dynamics, and ECM deformation. We use our model to quantify cell-cell communication mediated by ECM deformation and to show how this process depends on the mechanical properties of cells, the ECM fibres and the topology of the ECM network. In particular, we analyse the influence of ECM stiffness and cell contraction activity in the transmission of mechanical cues between cells and how the distinct timescales associated with different processes influence cell-ECM interaction. Our model simulations predict increased ECM deformation for stronger cell contraction and a sweet spot of ECM stiffness for the transmission of mechanical cues along its fibres. We also show how the network topology affects the ability of stiffer ECMs to transmit deformation and how it can induce cell detachment from the ECM. Finally, we demonstrate that integrating processes across different spatial and temporal scales is crucial for understanding how mechanical communication influences cell behaviour.

**Author summary:** The cell surrounding is a dynamic fibrous network known as the extracellular matrix (ECM). It supports and influences cell behaviour, playing a key role in cell communication, growth, and migration. Cells sense the ECM’s mechanical properties and exert traction forces on it, leading to the deformation of matrix fibres and the transmission of mechanical stress. These changes are transmitted along the ECM fibres, influencing the behaviour of neighbouring cells. Different subcellular structures and extracellular matrix components interact at various spatial and temporal scales, making mathematical modelling a valuable tool for analysing these interactions. We have developed a multiscale force-based model that quantifies mechanical stress transmission, captures cell detachment, and explores the impact of mechanical properties of both cells and the ECM. Our analysis shows that stronger cell contraction increases extracellular matrix deformation and suggests a range of extracellular matrix stiffness for effective mechanical cell-cell communication. We also use our model to investigate how ECM network topology can induce cell detachment by modifying the ability of stiff ECMs to transmit deformation when subject to cell-induced traction forces. Our results show the importance of coupling the processes occurring at different scales to capture the overall behaviour.

## 1 Introduction

Cells constantly interact mechanically with their microenviroment, notably defined by the structure and composition of the surrounding extracellular matrix (ECM). The ECM is a fibrous substrate composed of collagen, elastin, and other proteins, and forms a complex, interconnected and highly dynamic structure [1, 2]. The biochemical and biomechanical properties of this network regulate several cell processes, including spreading, growth, proliferation, migration, differentiation and morphogenesis [2–5]. The mechanical interactions of a cell with the fibres of the ECM network are mediated through the cell’s cytoskeleton, a viscoelastic and adaptive structure that enables cell movement and shape changes [6]. The dynamics of the cytoskeleton are regulated by active forces resulting from the polymerisation of its actin filaments [7], contractile forces produced by a class of molecular motors called myosins [8, 9] and interaction with the ECM. The traction forces produced by actomyosin activity are transmitted to the substrate through specialized connections known as focal adhesions (FAs) [10]. Located at the peripheral regions of the cytoskeleton, these structures connect the cytoskeleton to the ECM through clustered transmembrane receptors called integrins [11]. The regulation of the integrin binding and unbinding process within a FA plays a crucial role in both the outside-in and inside-out signalling [4, 12–16]. Regarding the former, integrins serve as biomechanical sensors as they respond to specific biochemical and physical characteristics of the cell microenvironment, initiating, upon stimulation, a cascade of events that lead to local changes in cytoskeletal dynamics. This results in the generation of mechanical forces, which causes overall changes in cell shape and motility. Over time, the response of integrins to the microenvironment influences transcriptional regulation, cell proliferation, differentiation, and survival. Moreover, integrins participate in signalling from the cell to the microenvironment, facilitating various cellular functions such as cell anchoring and the transmission of traction forces. This contributes to locomotion, substrate deformation, and ECM remodelling.

The ECM reorganises under cell-generated traction forces, leading to a strain-stiffening response [17, 18] and realignment of the ECM fibres [19–21]. The mechanical feedback between cells and the ECM have an important role in many biological processes. For example, in the process of vessel outgrowth (angiogenesis), the tip of nascent sprouts realign the ECM fibres along the direction of growing vessels [19]. The alignment of the matrix fibres then modulates the behaviour of the following endothelial cells to ensure vessel integrity and also guides vessel function (anastomosis) [22]. Abnormal changes in the ECM structure and composition are related to the initiation and progression of several cancers and other diseases [23, 24]. It has been reported that the invasive potential of cancer cells correlates with their increased ability to mechanically modify their microenvironment. In addition, it is known that mechanical stresses on the ECM surrounding primary tumours promote metastasis [25].

The mechanical interplay between cells and the ECM involves several complex and coordinated processes occurring simultaneously at distinct spatial scales. However, it is challenging to experimentally disentangle the temporal scales associated with the different processes of cell-ECM interactions, including cytoskeleton activity, FA formation and dynamics, and ECM deformation, and how they affect the overall dynamics. Mathematical modelling provides an alternative method for analysing the different temporal scales arising in this mechanical interplay. In addition, by using mathematical modelling, we can systematically investigate how variations in cell activity and biomechanical ECM properties affect the transmission of mechanical cues through the substrate, and, thus, mechanical cell communication.

In this study, we develop an agent-based model that simulates mechanical interactions between cells and ECM fibres and captures the simultaneous time evolution of the ECM, cells, and FA. We use the model to quantify cell-cell communication mediated by ECM deformation and show how this process depends on the mechanical properties of both the cells and ECM fibres and the topology of the ECM network.

Different modelling approaches have been proposed, focusing on particular aspects of this mechanical interplay: the transmission of mechanical cues through the ECM, the mechanosensitive response of the FAs to external forces and the internal cellular dynamics. We briefly review representative modelling works from each category:

### FA models

Previous modelling efforts [14, 26–28] have focused on understanding the dependence of FA dynamics on applied forces. Kong *et al*. [29] experimentally demonstrated that integrins within FAs form catch bonds with ECM fibres. These catch bonds explain the stick-slip behaviour of integrin clusters, which is characterised by an extended lifetime of integrin-ECM bonds at low cell-ECM interaction forces. However, as the forces increase, the lifetime of these bonds decays exponentially. Several modelling studies [14, 26, 27] have analysed force transmission through dynamic integrin-ECM bonds on the moving interface of the cell membrane. Novikova and Strom [14] further suggest that the fraction of bound integrins acts as a biological sensor of external ECM stiffness. Additionally, other studies [15, 30] have confirmed experimentally and theoretically that rigidity sensing emerges naturally from integrin dynamics, particularly in models incorporating catch bonds. In [28, 30], the authors investigate both experimentally and theoretically the role of talin and other FA-associated proteins in reinforcing the mechanical clutch, which regulates force transmission and converts mechanical stimuli into biochemical signals (mechanotransduction).

### ECM models

A wide range of mathematical models has been designed to capture and quantify the transmission of deformation, stress, and other mechanical signals through the ECM. Different modelling frameworks have been employed to describe ECM dynamics. For example, several modelling studies [21, 31–33] represent the ECM as an elastic network of interconnected fibres. There are other agent-based descriptions where fibres are represented by line segments that link and unlink from each other [34, 35]. Other works use continuum descriptions of the ECM stress tensor [36], the deformation gradient [21] and finite element descriptions of ECM fibres [20, 37] (see [38] for review). Regardless of the modelling approach employed, these models aimed to capture the stress distribution within the matrix [31, 33, 37], the realignment of fibres between two contractile cells [20, 31, 32], the impact of cell contraction force on the transmission of mechanical signals [33] and the influence of ECM network topology on the transmission of mechanical signals [31].

### Cell models

Several modelling approaches have been proposed to capture the dynamics of the cell internal structure [39, 40]. Whereas continuous representations of cells have been developed [36, 41], agent-based approaches are the frequent choice for capturing the effects of the internal activity and structure in individual cell behaviour. Examples of agent-based models of cells within a tissue include [32, 42–45] (see [46] for a recent systematic literature review). In a recent study [32], Tsingos *et al*. propose a hybrid representation, where cell structure is represented within the Cellular Potts framework and ECM fibres are represented as an elastic network. This approach captures the propagation of forces along the ECM, local cell-induced stiffening of the ECM and its deformation. In [42], Zheng *et al*. model cell migration within an elastic network of the ECM, using a centre-based model for cells. Although their model accounts for cell membrane protrusions and their attachment to the ECM, the dynamics of FAs are not included. Reinhardt & Gooch [43] represent cell structure by a collection of linear springs. This cell model is combined with an elastic model for the surrounding substrate which allows the authors to analyse the displacement of ECM fibres towards a contracting cell, the decay of their deformation with distance from the cell and ECM compaction in the region between two contracting cells. However, existing modelling works [32, 42, 43] have not considered the dynamics involving cell-ECM interactions, which include the formation of FAs and the evolution of their components.

Several theoretical models include the influence of the FA dynamics on cell behaviour. For instance, recent studies [44, 45] combine the active dynamics of the cell cytoskeleton with a mechanosensitive response of the adhesions to an elastic substrate. This approach allows the authors to reproduce active cell processes involved in the cell migration cycle, such as the formation of FAs, generation of protrusions due to actin polymerisation, and active contraction.

The coupling across different spatial and temporal scales remains a challenge from the theoretical perspective [38]. A few modelling studies which aim at capturing cell-ECM interactions across different scales include a recent model by Keijzer *et al*. [47]. They extend the model introduced in [32] by introducing FA mediated adhesions.

As previous modelling efforts do not account for the force-based effects of the FA dynamics on cell-ECM interactions and mechanical cell communication, we propose a new multiscale force-driven model that combines three building blocks: (*i*) an elastic model describing the ECM fibre network, (*ii*) a cell model accounting for its structural and active forces, and (*iii*) cell-ECM interactions incorporated through the formation and dynamics of FAs, which are described by the mechanosensitive binding-unbinding processes of integrin clusters. Integrating the processes involved in the cell-ECM interaction provides a holistic understanding of the interaction between these three elements, considering the different timescales involved. We also introduce a number of metrics that permit us to quantify the local ECM orientation, the anisotropy of its fibres and to measure transmission of mechanical cues through the ECM. We examine how the mechanical properties and ECM network topology influence the ECM deformation capabilities and the rate of stress transmission and realignment within is fibres. We can also examine how overall behaviour is determined by the balance in the transmission of mechanical cues through the ECM and the integrin dynamics within the regulation of FAs.

We use our model to investigate cell-ECM interactions, and the crosstalk between cells via mechanical cues in different scenarios. We begin by illustrating the dynamics of FAs in a single round contracting cell placed on an ECM substrate. We then consider a scenario where two contracting elongated cells are in mutual proximity within the ECM and quantify the influence of key properties of the system, such as ECM stiffness and active contraction force, on the transmission of mechanical cues. In particular, our model simulations predict increased ECM deformation for stronger contraction activity and a sweet spot of the ECM stiffness for the transmission of mechanical cues between cells. Moreover, we study how the local ECM topology affects its *effective* rigidity, i.e. its capability to transmit deformation under traction forces, and induce cell detachment. Our model formulation also permits us to analyse the distinct timescales associated with different processes involved in the cell-ECM mechanical feedback, clarifying their influence on the resulting behaviour.

The remainder of this paper is organised as follows. In section 2 we describe the different building blocks of our multiscale model, and present some spatial metrics that permit us to quantify the realignment of the ECM fibres and stress transmission along its fibres. In this section we also include a model simulation that illustrates the dynamics of FAs in a single contracting cell placed on an ECM substrate, and a brief analysis of the FA model, i.e., the cell-ECM interaction model. In section 3, we consider a scenario of two contracting elongated cells placed in mutual proximity within the ECM. By using the metrics defined in section 2, we perform numerical simulations of our model to conduct an extensive parameter sensitivity analysis and quantify variations in the system behaviour induced by changes in ECM stiffness and active contraction force. We then further analyse the effects of the local ECM topology on the transmission of mechanical cues and cell detachment. In section 4 we present a summary of the main conclusions of our work and future perspectives.

## 2 Materials & Methods

This section is organized as follows: We begin in section 2.1 with a general overview of the model, where we describe the various components involved in the interaction between cells and the ECM. Next, we introduce the metrics used to quantify the alignment of ECM fibres (section 2.2) and the stress transmission between cells via the ECM (section 2.3). We then present in section 2.4 an illustrative simulation of a single cell contracting, which demonstrates how the various components of the model are interconnected and evolve simultaneously. Finally, in section 2.5, we analyze the FA model, which describes the interaction between cells and the ECM. This includes a bifurcation analysis that illustrates how the forces exerted on the FA can lead to cell detachment from the ECM.

### 2.1 General overview of the model

We formulate a multiscale model of the mechanical interactions between cells and the extracellular matrix (ECM). The model comprises three building blocks. Firstly, an agent-based model captures cell shape changes caused by the mechanical stimuli from the ECM. Secondly, the ECM is modelled as an elastic material composed of elastic fibres that are interconnected by crosslinks. Finally, the cells and ECM dynamics are coupled via integrins. Integrins are cross-membrane receptors that cluster within FAs and bind ligands embedded within the ECM. Integrins are mechanosensitive and play a crucial role in transmitting mechanical stimuli from the ECM to cells. Figs 1A-C illustrate the main components of our model and their interactions. We distinguish each fibre within the ECM, each cell and the FAs that mediate their interactions. Our model is force-based, i.e. the dynamics of the different components are driven by balancing the mechanical forces that act upon them. A schematic of these forces is presented in Fig 2.

**Fig 1.**
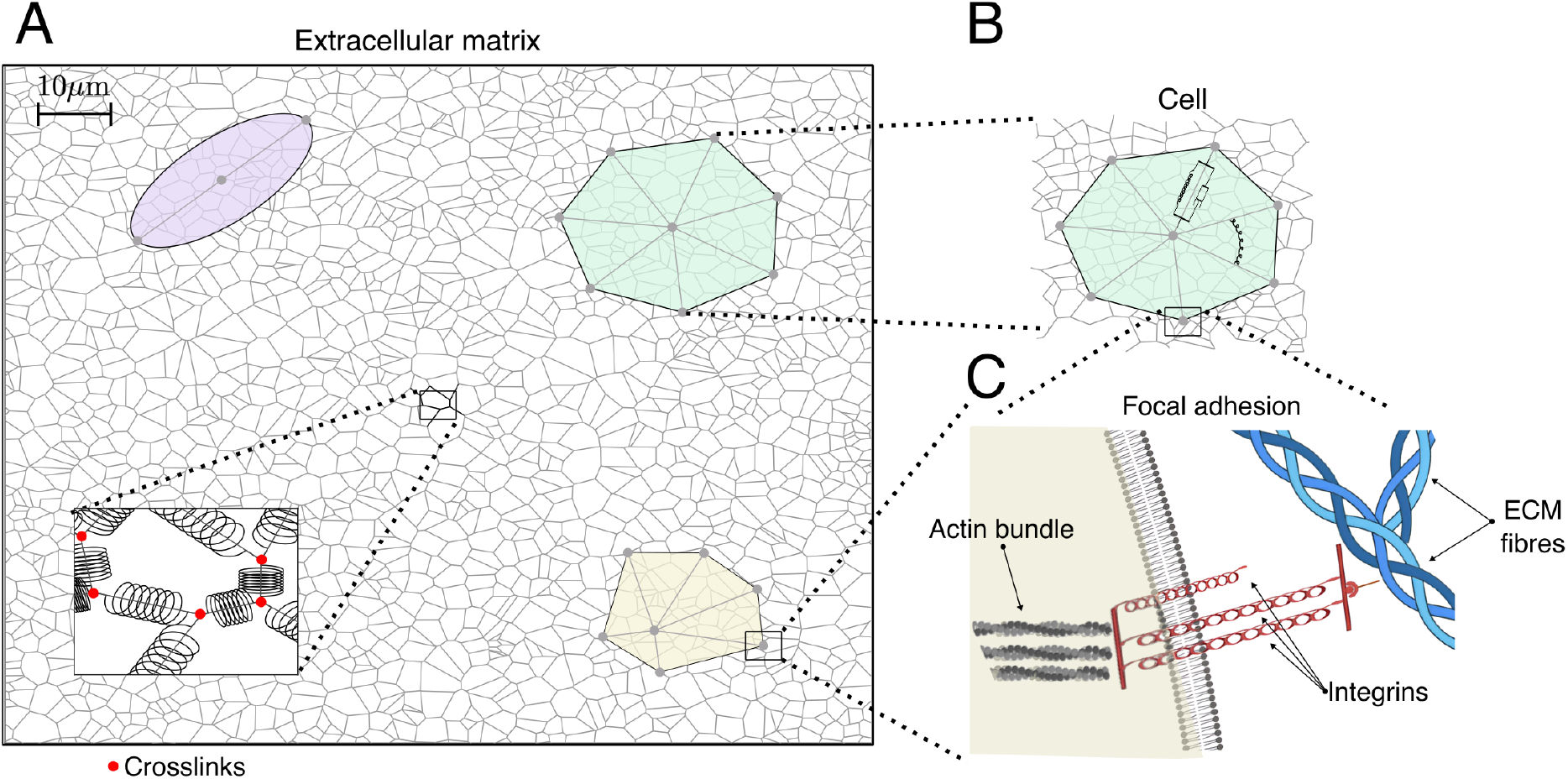
Schematic representation of the multiscale model. **(A)** A cartoon illustrating the ECM network. Following [31], given the density of ECM fibres, we randomly distribute a set of points which are the seed of our ECM configuration. Then, the ECM network is generated by constructing a Voronoi triangulation on the seed points. The set of edges in this construction are the ECM fibres. We also illustrate different cell geometries: an elongated cell (purple), a round spread-out cell (green) and a fan-shaped cell (yellow). **(B)** Cells are represented as irregular polygons. Their vertices are given by the ends of viscoelastic segments that represent actin bundles that originate from the organising centre of the cell cytoskeleton (which does not necessarily coincide with the polygon centre of mass). These bundles are modelled as viscoelastic elements that deform when subject to forces exerted by the ECM which are transmitted by the integrins within the FAs. Cells can also contract under the action of myosin motors. Angular forces between these segments enable cells to maintain their geometry. The polygon vertices represent the sites where FAs are formed. **(C)** A cartoon showing our representation of FAs in which integrins, trans-membrane receptors which bind ECM fibres, are modelled as clusters of linear springs in parallel.

**Fig 2.**
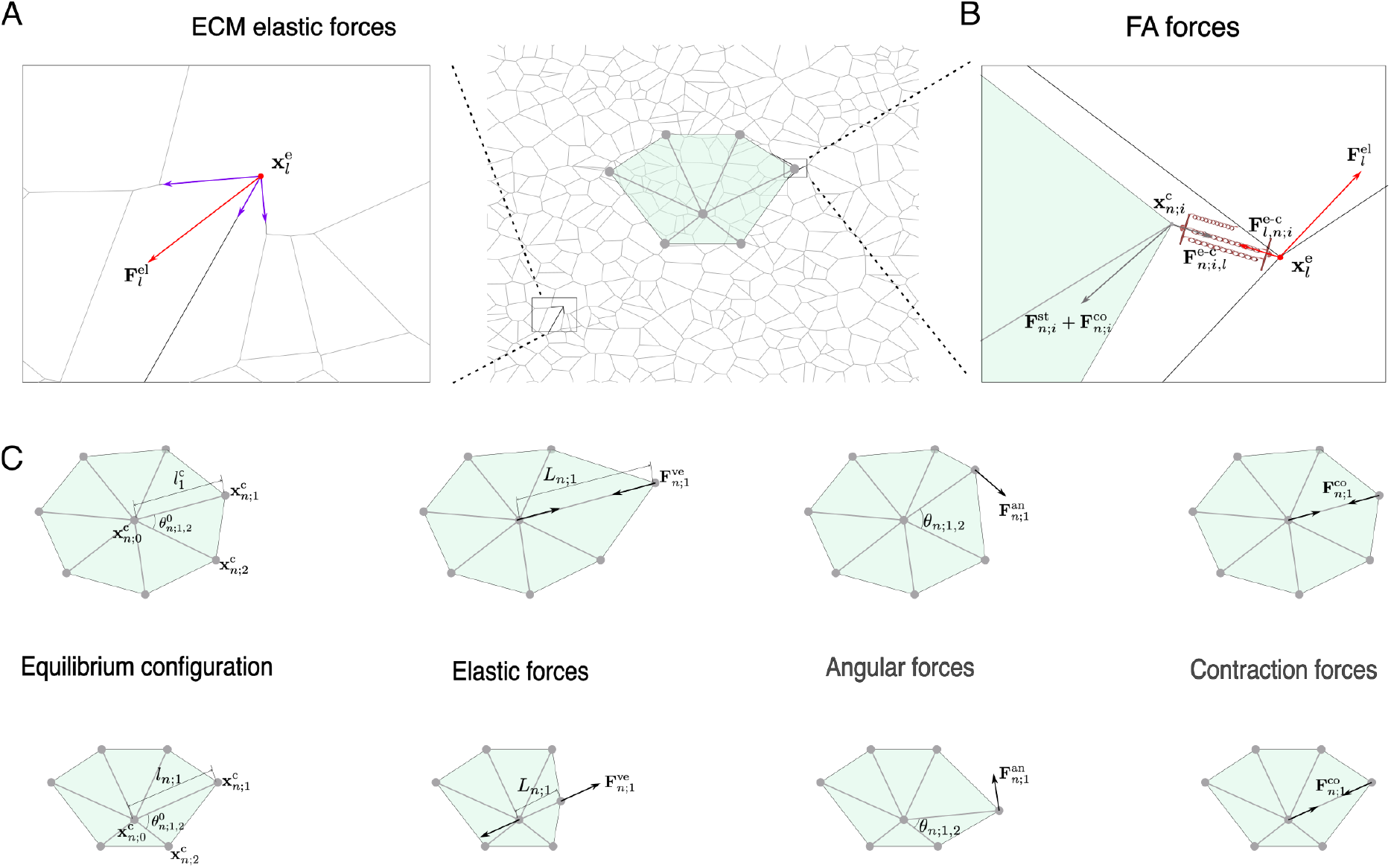
A schematic representation of the forces included in the model. **(A)** A zoom view of a randomly perturbed ECM region. The resultant force acting on the *l*-th crosslink is shown in red, and the elastic forces exerted by individual fibres at this crosslink are highlighted in purple. **(B)** A close-up view of the mechanical forces acting on an FA. Here, ECM (cellular) forces are shown in red (grey). In addition, we illustrate the cell-ECM interaction force, 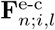, acting between the ECM crosslink (in red) and the cell binding site (grey). **(C)** An illustration of the different structural and active forces incorporated into our cell model. From left to right: equilibrium cell configuration, elastic forces, angular forces and active contraction forces. Two different cell geometries are shown as representative examples: a round (at the top) and a fan-shaped cell (at the bottom).

#### Model variables

Before detailing the various components of our model, we clarify the notation used, particularly we list the model’s elements and how we apply subindex and superindex notation to represent them. In our model simulations, we track the positions of ECM crosslinks, 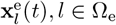, positions at which individual matrix fibres are bound together, Ω_e_ is the set of crosslinks. The number of cells in our model is given by *N*_c_, and each cell is represented by the index *n*. The cells are defined by their organising centre, whose position is given by 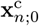, and set of vertices, whose positions are denoted by 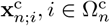. where the set of vertices of a certain cell *n* is 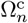. The number of vertices that define a cell is assumed constant and is given by 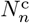. Each cell vertex represents a cell adhesion site (location where a FA is formed, i.e. interacting points). We also record the evolution of the number of bound integrins 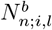 that form a FA and connect the cell vertex 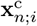 with a nearby ECM crosslink, 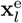. We list the variables of the system in Table 1.

**Table 1.**
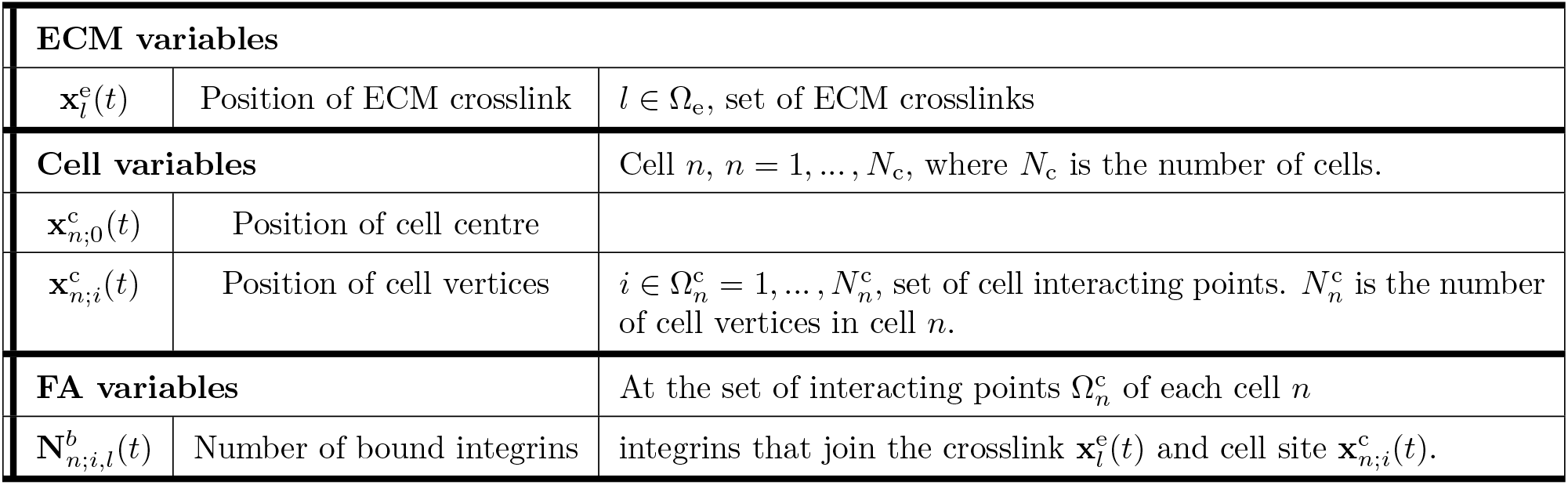
Variables of the model.

#### ECM model

We view the ECM as a network whose edges represent individual fibres and whose nodes represent crosslinks, i.e. sites at which individual matrix fibres are bound together. These bonds provide structural stability to the ECM network [48]. The position of each crosslink is referred to as 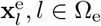, where Ω_e_ is the set of all crosslinks in the network, see Table 1. Following [31], we assume that the fibres are elastic and thus can be described by linear springs (see Fig 2A). In the absence of external forces, a crosslink 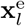 evolves from an arbitrary initial position to an equilibrium configuration where all the elastic forces are in equilibrium. In addition to these *internal* forces, the ECM is also subjected to forces applied at specific points within the ECM (see Fig 2B). These points correspond to where cells have formed FAs that enable them to bind to the ECM. Active cellular behaviours, such as contraction driven by myosin motors, give rise to *external* forces which act upon the ECM and alter its (equilibrium) configuration.

In the overdamped limit, the evolution of crosslink positions is determined by balancing the (internal) elastic forces acting between neighbouring crosslinks and the (external) forces due to active cellular processes:

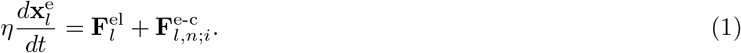

In (1), 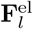 is the resultant elastic force acting on the *l*-th crosslink (see section Detailed model description in S1 File) and 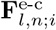 represents the interaction force between this crosslink and *i*-th vertex of the *n*-th cell (see below, in the description of the mechanical coupling between the cells and the ECM). The parameter *η* denotes the damping coefficient.

#### Cell model

Cells are viewed as agents with polygonal shapes (see Fig. 1A). In our model, we define *N*_c_ different cells. Each cell *n, n* ∈ 1, …, *N*_c_, is defined by a set of points, 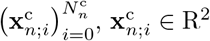. We denote by 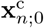 the position of the cytoskeleton organising centre. The geometry of the cell is then modelled by a set of viscoelastic segments that emanate from the organising centre and terminate at the cell vertices, 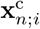. Here, 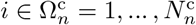, with 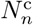 being the total number of structural segments in cell *n* (see Table 1 for details of the notation). 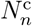 can be varied to accommodate different cell shapes. Biologically, these segments can be understood as bundles of actin filaments or stress fibres that provide structural support to the cellular geometry [40, 49]. They also correspond to the sites of cell-ECM interactions mediated by FAs [50]. In Fig 1A, we illustrate three possible cell geometries corresponding to an elongated (top-left), a round (top-right) and a polarised fan-shaped (bottom-right) cell.

The forces that drive the cellular dynamics can be divided into three classes (see Fig 2B-C):

- **Internal structural forces**, 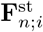 These forces regulate cell shape. We consider two types of structural forces. First we account for the viscoelastic properties of actin bundles contained in the stress fibres. We describe the viscoelastic behaviour of the structural segments of the cell by the Kelvin-Voigt model [40, 49]. We also account for so-called *angular forces*, which act to maintain cell shape. These angular forces set a target or natural angle between consecutive stress fibres, just as elastic springs have a natural length. When the angle deviates from its target value, a restoring force acts to restore the angle to its target value, and prevent the cell from collapsing upon itself.
- **Active contraction forces**, 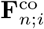. Active forces are generated within cells and usually driven by
- **out-of-equilibrium ATP-dependent reactions.** They are essential for most cell functions, including locomotion and adhesion. The present model includes only contraction forces. These forces are propagated along the actin bundles and are generated by myosin motors, which produce contractile forces on actin filaments [51]. Specifically, we represent active contraction forces as constant forces that contract the viscoelastic elements towards the cytoskeleton organising centre.
- **Mechanical interactions with ECM**, 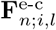. These interactions are mediated by FAs, specialised anchoring points at which cells bind to the ECM. A detailed description is given below, in the paragraph describing the mechanical coupling between the cells and the ECM.

A more detailed description of the forces is given in section Detailed model description in S1 File. In our model we assume that cells are sparsely seeded on the ECM, so direct cell-cell interactions are negligible. In the overdamped limit, where the inertial effects can be ignored relative to the viscous forces, the positions of the cell centre and vertices of the *n*-th cell are given by:

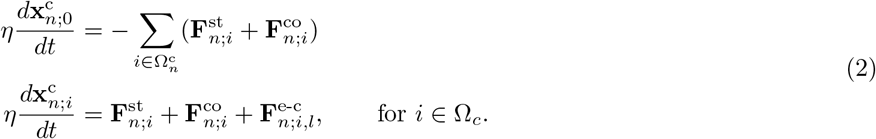

The solution of Eq (2) determines the evolution of the cell shape.

#### Mechanical coupling between the cells and the ECM

FAs are complex macromolecular clusters that transmit mechanical forces and regulatory signals between the ECM and an interacting adherent cell [11, 12]. The formation of FAs is a complex process which involves inside-out cell mechanisms (signals originating from within the cell change the behaviour of surface receptors, affecting their ability to interact with external molecules), in conjugation with outside-in mechanisms (ECM biochemical, biomechanical composition and forces regulate the composition of FAs) [52, 53]. In addition to anchoring cells to the ECM, FAs act as hubs for transmitting information from the ECM to the cells. This process involves hundreds of proteins, many of which associate and dissociate in a dynamic manner [13]. Integrins are the main component of FAs. They are the transmembrane mechanosensitive receptors that create mechanical bonds between the actin cytoskeleton and ECM fibres by binding to ligands embedded within the latter [13]. When subjected to mechanical stress, integrins deform and transmit mechanical stimuli from the ECM to the cell. Integrins are normally modelled as linear springs, i.e. they are assumed to stretch at a rate proportional to the applied force [14, 26–28]. Furthermore, the mechanical state of integrins also affects their binding kinetics [29, 30]. This aspect of integrin mechanosensitivity has previously been modelled by assuming that the unbinding rate, K_off_, is a function of the force applied to the integrin [14, 30].

In our model, cell vertices represent the ends of actin bundles, which correspond to the sites of FA assembly. Integrins within FAs transduce mechanical signals between cells and the ECM. They can be deformed due to the forces exerted by the ECM. This deformation, in turn, exerts forces on the actin bundles which, among other effects, can change the cell geometry. Similarly, forces exerted by the cell (e.g. myosin-motor mediated contraction) are transmitted to the ECM by stretching the integrins which, in turn, deforms the ECM fibres locally. For simplicity, we assume that integrins within FAs are organised as arrays of parallel linear springs (see Fig 1C). Therefore, all integrins undergo the same amount of deformation and the force exerted by the array of integrins is proportional to the number of bound receptors. This implies that to characterise the force exerted by the set of integrins, we must determine both their deformation (stretch) and the number of integrins bound to the ECM.

We assume that the cell-ECM mechanical linkages connect a cell vertex, 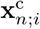, and a matrix crosslink, 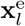 (see Fig.1C). We denote by 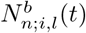 the number of integrins bound to the ECM, 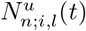 denotes the number of unbound integrins at time *t*. Following [14], we assume that the total number of available integrins at a FA is conserved

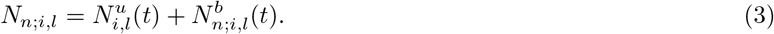

Based on our assumption that integrins behave as an assembly of parallel Hookean springs, the resulting force exerted by a cluster of bound integrins, 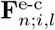, is determined by:

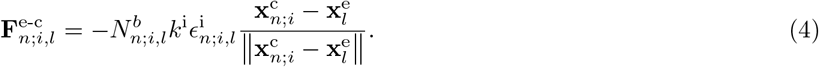

In (4), *k*^i^ denotes the integrins elastic constant and 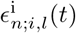 defines the extension of the integrins, which is given by:

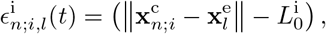

where 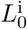 is the equilibrium length of an integrin.

The net force exerted by the bound integrins, 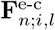, results from their deformation within a FA (see Fig 2B). This deformation is generated by the forces exerted by the ECM (elastic forces exerted by fibres converging at 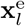) and the forces exerted by the cell (structural and active forces at 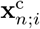). By Newton’s third law, we know that the forces exerted at the cell adhesion site, 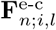, and the ECM crosslink, 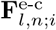, have equivalent magnitude but pointing in opposite directions: 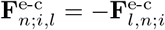.

Lastly, following [14, 26], we assume that integrins within FAs bind and unbind at rates which depend on the mechanical forces caused by their deformation, 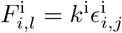. Specifically, we use the so-called catch model [14, 29] which has been proposed to more accurately describe the detachment of integrins. This model assumes that the maximal lifetime of a cell-ECM bond occurs at intermediate forces, whereas for large forces, it decays exponentially. This is captured by assuming that the integrin unbinding rate, 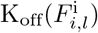, depends on the force which results in its deformation. The total number of available integrins in a FA, *N*_*n*;*i,l*_, and their binding rate, K_on_, are assumed to be constant. Combining these effects, we arrive at the ODE for the number of bound integrins in a FA, which together with (3):

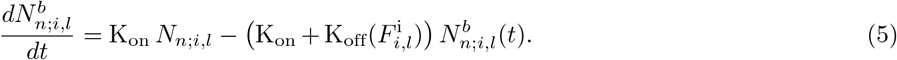

To summarise, our model accounts for cell-ECM dynamics that span different spatial scales, which include ECM deformation, sub-cellular processes (such as myosin-driven contraction) and integrin binding and unbinding processes. In particular, the positions of ECM crosslinks are determined by Eq (1), and the deformation of each cell is governed by Eq (2), with cell-ECM coupling captured by the formation of the FAs. The number of bound integrins at each FA is given by Eq (5). A more detailed description of the model can be found in section Detailed model description in S1 File.

We seek to quantify the effects of cell contraction on the ECM, i.e. to determine the deformation of the ECM caused by pulling forces due to cellular activity. We introduce several metrics to quantify fibre alignment and the transmission of mechanical stress between contracting cells.

### 2.2 Measuring fibre alignment

#### Local alignment metric

To quantify the effects of active cell processes (such as myosin-driven contraction) on ECM structure, we introduce an alignment metric. Following [31], we consider a local orientation tensor **Θ**_*l*_ which is defined at each ECM crosslink and accounts for the degree of alignment of the fibres converging to it. At crosslink *l* we have:

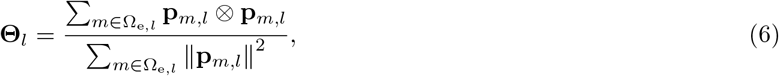

where ⊗ is the tensor product, the sum is over the set Ω_e,*l*_ of neighbouring nodes of a crosslink *l*, and

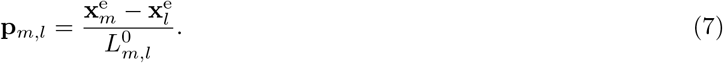

If the set of vectors {**p**_*m,l*_, *m* ∈ Ω_e,*l*_} that converge at crosslink *l* contains at least two non-parallel vectors, then the local orientation tensor, **Θ**_*l*_, is symmetric and positive definite with Tr(Θ) = 1. As such it can be diagonalised and its eigenvalues {*λ*_*l*;1_, *λ*_*l*;2_} lie within the interval (0, 1). When the fibres are arranged isotropically, **Θ**_*l*_ has two identical eigenvalues *λ*_*l*;1_ = *λ*_*l*;2_. By contrast, if the fibres are distributed anisotropically, then the eigenvalues are distinct *λ*_*l*;1_≠ *λ*_*l*;2_. We note also that the eigenvector corresponding to the largest eigenvalue is parallel to the direction of the preferential orientation of the ECM fibres at this crosslink. The local orientation 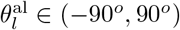 points in the direction associated with the largest eigenvalue. The anisotropy metric, 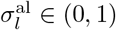, describes the local anisotropy of ECM fibre distribution at the *l*-th crosslink:

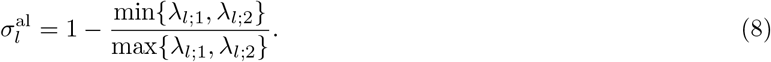

In near-to-isotropic scenarios, 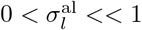, while for anisotropic cases (i.e. fibres converging at crosslink *l* are strongly aligned in the same direction) 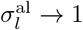. Fig 3A illustrates the principal alignment directions and the corresponding anisotropy parameter for three cases of increasing anisotropy. The left panel corresponds to the isotropic case, while the right panel corresponds to strong vertical alignment of ECM fibres.

**Fig 3.**
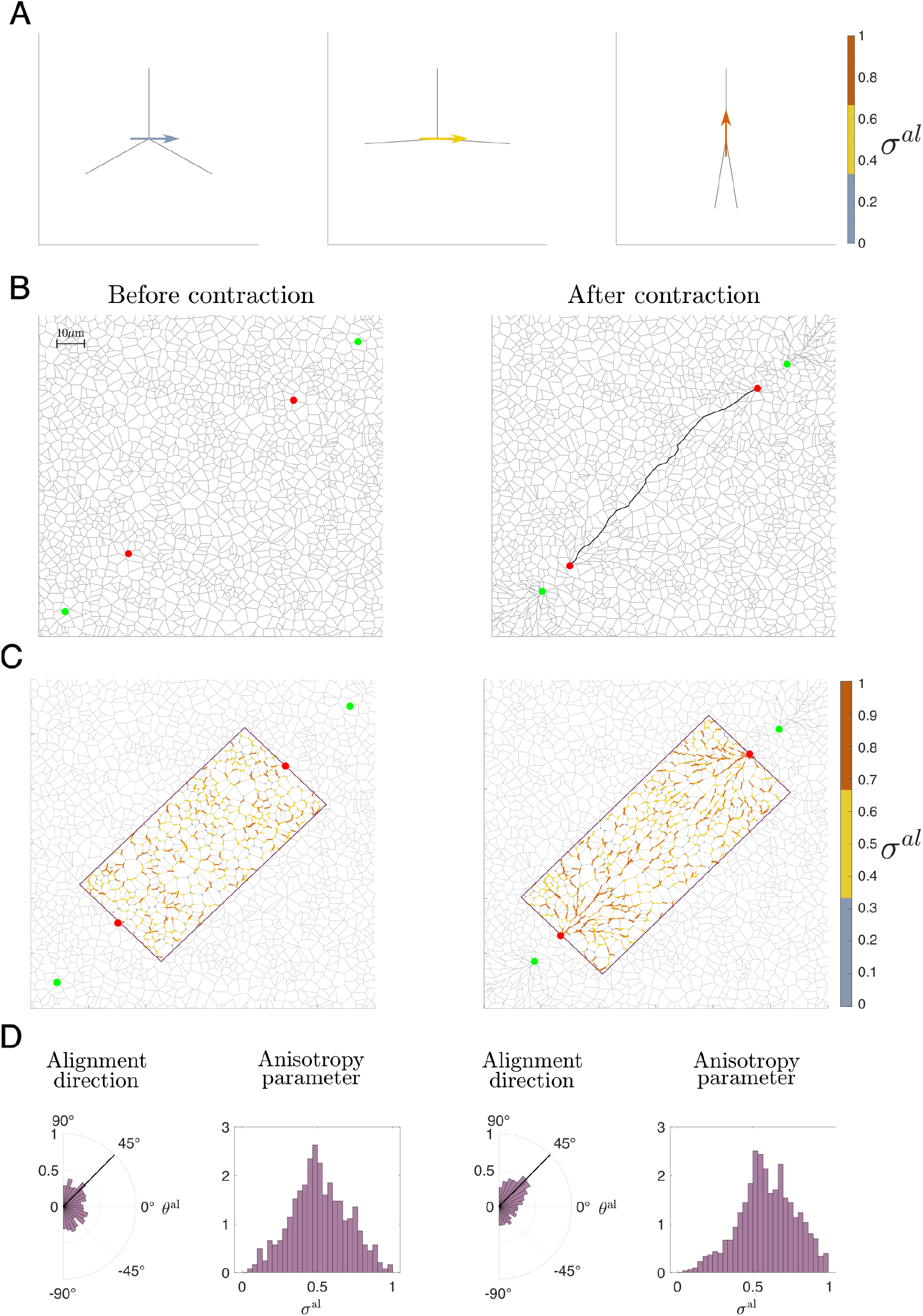
An illustration of the alignment measure. **(A)** Examples of local alignment measures *σ*^al^ (see Eq (8)) for isotropic (left), partially anisotropic (middle) and strongly aligned (right) fibres. Here, ECM fibres are shown in light grey. The alignment vector corresponding to the principal eigenvalue of the tensor, **Θ**^*µν*^, from Eq (6) is coloured according to the value of the anisotropy parameter, *σ*^al^. Blue corresponds to isotropic fibre distribution 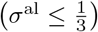, yellow indicates intermediate alignment 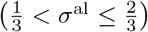, and orange corresponds to an anisotropic case 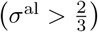. **(B)** ECM configurations at equilibrium (left) and after the contraction of two pairs of crosslinks (coloured in red and green), corresponding to two cells. The maximum stress transmission path after cellular contraction is highlighted in black. **(C)** Local alignment measures in the ECM fibres at an equilibrium (left) and after cellular contraction. ECM fibres within the region between the two cells (outlined in purple) are coloured according to the corresponding value of the local fibre anysotropy parameter *σ*^al^. **(D)** Histograms of distributions of the principal eigenvector and the anisotropy parameter in the purple regions for the ECM configurations shown in **(C)**. After cell contraction, ECM fibres realign along the contraction direction (≈ 45^*o*^), and the distribution of the anisotropy parameter skews towards an anisotropic scenario.

#### Propagation of ECM alignment induced by active cell contraction

We determine how fibre alignment is transmitted between two contracting cells by calculating the alignment direction,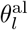, and the anisotropy parameter, 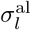, for ECM fibres between two contracting cells (outlined in purple in Fig 3C). We then plot the distribution of these metrics in the ECM region of interest before and after cellular contraction (see Fig 3D). For the relaxed scenario (Fig 3D, left panel), we note the uniform distribution of the local fibre orientation and the symmetric distribution of the anisotropy parameter around a partially aligned state 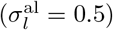. After 5 s of cell contraction (Fig 3D, right panel), the ECM fibres align in the direction of the vector connecting the two cells, causing the anisotropy parameter distribution to skew towards an anisotropic scenario.

To make this comparison quantitative we analyse the distribution of the anisotropy parameter before and after cellular contraction. Specifically, we calculate the mean and skewness of the alignment metric distribution. The latter allows us to quantify the shift of the alignment metric towards larger values when active cell contraction affects the ECM.

### 2.3 Percolation: measuring the spatial stress transmission

Another crucial aspect of mechanically-mediated cell-cell communication is the spatial propagation of stress through the ECM fibres. To quantify this effect, we propose to measure the stress percolation between ECM crosslinks. This technique involves determining a stress transmission path in the following way:

1. We calculate the tensional stress, 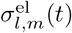, of the stretched fibres that comprise the ECM after their deformation due to cell contraction, i.e. those fibres whose length at time *t* is greater than their length at *t* = 0:

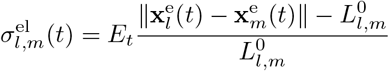

for a pair of crosslinks *l, m* whose separation at time *t* is greater than their initial distance 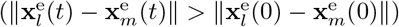.
2. We determine the set of fibres, Ω_Path_, that compose the path connecting two crosslinks within the ECM 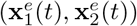 as those that maximise the stress transmission. The crosslinks with positions 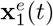 and 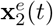 correspond to the two points of cell-ECM interaction (see Fig 3B). To find a path whose total stress is maximal, we utilise the Dijkstra algorithm [54], minimizing 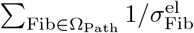. Where Fib = *l, m* refers to the pair of crosslinks that form a fibre. If there is no such connected path, we conclude that there is no mechanical communication between the cells.
3. We then calculate the total stress 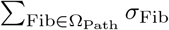 in all fibres in Ω_Path_. To quantify the degree of tortuosity of this path, we compare the Euclidean distance between the nodes with the length of the path: 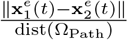 where dist(Ω_Path_) is the sum of lengths of the fibres that form the path. When the tortuosity is closer to 1, there is more deformation in the path connecting both cells; when it is small, there is less transmission of mechanical cues.

In Fig 3B (right panel), we illustrate the percolation path between two ECM crosslinks (indicated by red points) after the contraction between the two pairs formed by a red and a green crosslink.

### 2.4 Focal adhesion dynamics for an isolated cell

To test the model’s ability to capture dynamic interactions between ECM crosslinks and FAs of a round contracting cell, we simulate an isolated cell attached to a randomly oriented (isotropic) ECM. Fig 4A-B show the initial and final configurations of the cell-ECM system after 5 s of constant cell contraction. The initial configuration is obtained by letting the system relax to its equilibrium configuration, with the cell attached to the ECM and no contraction activity, i.e. we simulate the system of equations setting *F* ^co^ = 0. Once the system reaches the latter state, we use this configuration as an initial condition for the model. Then the cell contracts under the action of active contractile forces, i.e. *F* ^co^≠ 0. As expected, the transmission of stress attenuates with distance: from the adhesion sites the stress emerges in radial paths, with compressed transversal fibres along these paths.

**Fig 4.**
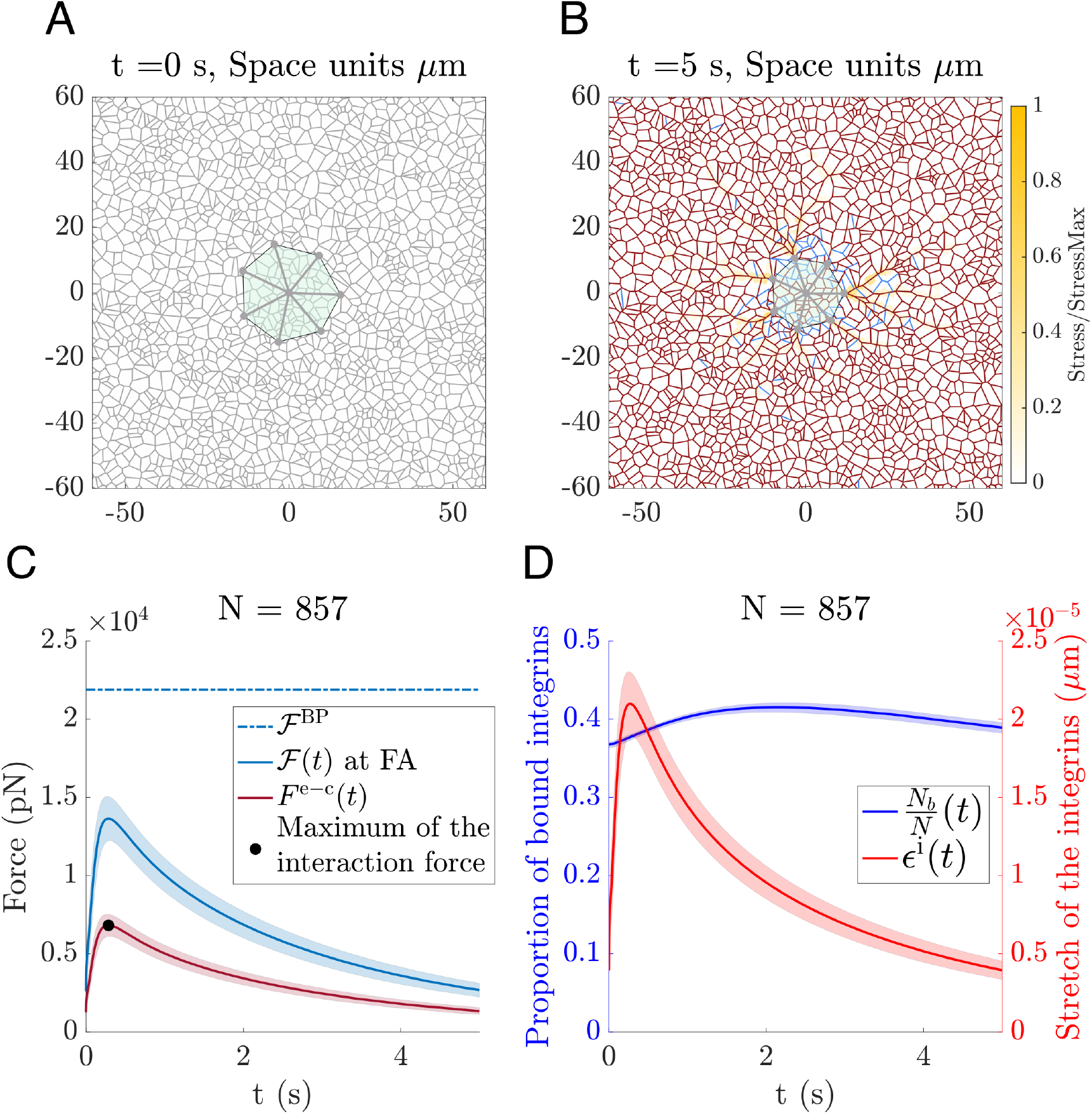
Single cell contraction. **(A)** The initial configuration of the cell-ECM system before cell contraction. A round cell is placed on top of a relaxed ECM network (as described in section Detailed model description in S1 File). The fibres in equilibrium are shown in grey. **(B)** ECM and cell configuration at *t* = 5 s after contraction. Stretched fibres are plotted in dark red, and the compressed fibres are in blue. The colour map indicates the stress of the ECM normalized by the maximum stress in the ECM. Parameters: Young’s modulus of the ECM fibres *E*_t_ = 10MPa, number of integrins per FA *N* = 857, *F* ^co^ = 20 nN. The remaining parameters are listed in S1 Table. **(C)** The time evolution of the mean force exerted at the FA (solid blue line) and the elastic force exerted by the FA (dark red line) for the simulation from **(A)** and **(B)**. The bifurcation force of the FA for this system is shown by a dash dotted blue line. The black dot indicates the peak tension force exerted by the FA. Shaded regions indicate the standard deviation calculated over all the cell FAs. **(D)** Evolution of the proportion of bounded integrins per FA (in blue) and the stretch of the integrins (in red) during cell contraction.

A key feature of our model is that it seamlessly accommodates the different time scales involved in the cell-ECM system. In particular, the model captures the evolution of the different forces involved in the cell-ECM system.

Integrin deformation and detachment is greatly affected by the forces applied to the FA due to cell contraction and ECM deformation, i.e. the contractile mechanisms and the viscoelastic properties of both the cell and the ECM determine the dynamics of their interaction (see section Detailed model description in S1 File). In the specific simulation shown in Fig 4, the model shows the tension undergone by the FA peaking early to a value below the detachment threshold. The FA tension then relaxes as the transmission of the deformation through the ECM progresses (see Fig 4C).

Finally, the evolution of the focal adhesion components, specifically the ratio of attached integrins and their stretch, are illustrated in Fig 4D. When analysing the temporal evolution of the focal adhesion components, we found that the dynamics of the interaction forces are mainly influenced by the stretching process of the integrins: while the interaction forces and stretch of the integrins in Fig 4B-C have similar timescales, the changes in the number of bound integrins occur on a longer time scale. As predicted by the model analysis presented in section 2.5, the binding-unbinding dynamics are slower than those associated with integrin deformation.

### 2.5 Analysis of the FA model

To gain further insight into the FA dynamics, we performed a bifurcation analysis of the force-binding model by varying the strength of the force applied to the FA system, which we call external force in this section. For simplicity, we focus on the dynamics of a specific FA, and modify our notation as follows: *N*_*b*_ and *ϵ*_*b*_ refer to the number of bound integrins in the FA and their stretch respectively, *N* is the total number of available integrins and 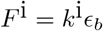 the force resulting from deforming a single integrin.

With this notation, the system of ODEs for the FA model is as follows:

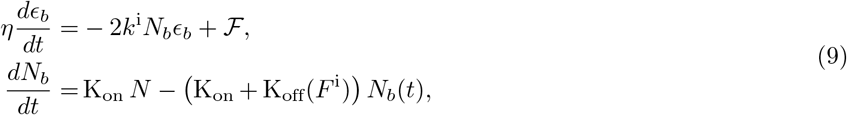

where ℱ denotes the external force applied to the assembly of integrins, i.e. forces resulting from cell activity and ECM deformation. We performed a bifurcation analysis of Eq (9), viewing the force, ℱ, as a control parameter.

Fig 5A shows that the system exhibits a saddle-node bifurcation. For values of ℱ smaller than a critical value, *F* ^BP^, the system exhibits monostability. In particular, when ℱ *< F* ^BP^, a positive equilibrium, which corresponds to a stable cell-ECM bond, coexists with an unstable equilibrium. For ℱ *> F* ^BP^, the system evolves to a state in which no bound receptors exist, which we interpret as detachment from the ECM. This is illustrated in Fig 5C, where we plot trajectories corresponding to attachment or detachment.

**Fig 5.**
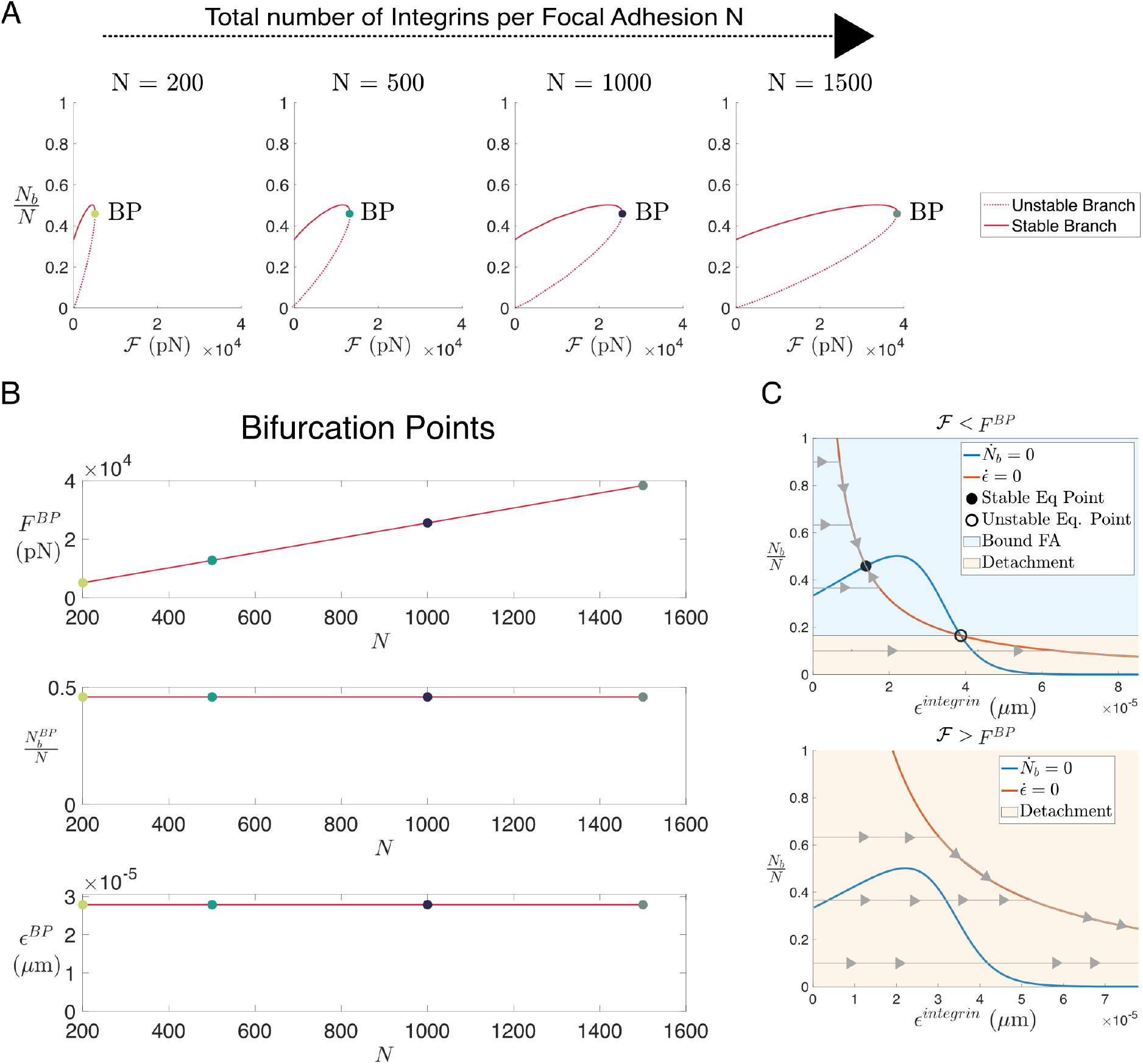
Bifurcation analysis of the FA ODE system. **(A)** A series of plots showing bifurcation diagrams of the system described by Eq (9) as the force parameter, ℱ, is varied. The bifurcation diagram is obtained numerically using BifurcationKit from Julia [55]. The four panels from left to right correspond to the increasing total number of available integrins *N* = 200, 500, 1000 and 1500. **(B)** Three plots showing the values of the bifurcation force, the ratio of attached integrins in the FA, and their stretch at the bifurcation point for different values of the total available integrins, *N*. Coloured circles represent the values obtained numerically as in **(A)**; red curves are calculated from an analytical expression (see section Exact calculation of the critical value *F* ^BP^ in S1 File). **(C)** Phase portrait diagrams for the case of *N* = 1500 for a value of the force ℱ lower than the bifurcation value *F* ^BP^ (top) and a value higher than the bifurcation value (bottom). In grey, we show trajectories of the solution of Eq (9) starting from the initial condition of zero-stretch and different proportions of bound receptors, *N*_*b*_*/N*. The nullclines of the system are shown in orange for *dϵ*_*b*_*/dt* = 0 and in blue for *dN*_*b*_*/dt* = 0. The region shaded in blue represents the basin of attraction of the stable fixed point, while the trajectories that enter the copper region will lead to the detachment of the cell from the ECM since the number of bound receptors tends to zero.

We also analysed the system behaviour as the number of receptors, *N*, varies. Fig 5A shows that as *N* increases, the size of the region of stability also increases (*F* ^BP^ increases as *N* grows), which means that ECM binding becomes more robust when integrins are more abundant. In Fig 5B, we show how the system variables at the bifurcation point depend on the total number of available integrins *N*. We note that our model predicts a linear relationship between the number of available integrins and the bifurcation force, *F* ^BP^. However the proportion of bound integrins and their stretch remain constant at the bifurcation point, regardless of the changes in the total number of available integrins *N*. We conclude that the model exhibits a scale invariance property as the number of available integrins *N* changes: the proportion of bound integrins and the ratio of the force exerted per available integrin do not depend on the number of available integrins (see Fig 5A,B). We analyse the system dynamics in terms of these ratios by dividing the system given by Eq (9) by *N*. Although the equilibrium points are not altered, the dynamics of the stretch changes; for large values of *N*, the stretch dynamics occur much faster than in cases with low values of *N*, indicating that a quasi-equilibrium approximation may be appropriate in this scenario.

In section Exact calculation of the critical value *F* ^BP^ in S1 File we derive an implicit expression for the bifurcation force *F* ^BP^, and related expressions for the proportion of bound integrins 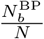 and their stretch 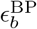:

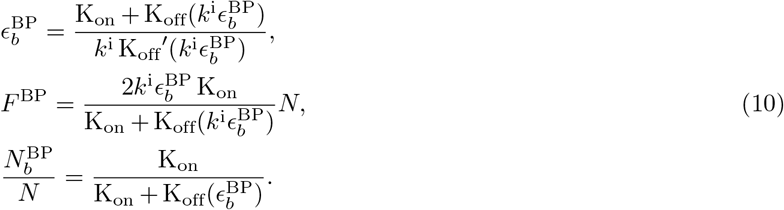

The scale invariance property is reflected in the bifurcation values, as 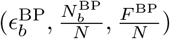 only depend on the binding-unbinding ratio.

These results show how forces influence cell-ECM interactions. In our model, the forces acting on each FA arise from a combination of viscoelastic forces, generated by both the cells and the ECM, and active cell contraction. For the interaction between the cell site 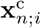 and the ECM crosslink 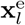, the force is therefore given by

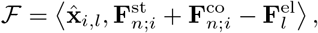

where ⟨, ⟩ refers to the inner product and

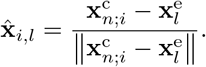

In the next section, we investigate how active and viscoelastic forces are transmitted through FAs.

## 3 Results

### 3.1 Reorientation and deformation of the ECM influence cell-cell interactions at a distance

Fig 6 shows simulation results for two elongated contracting cells located within the ECM at a distance from each other. These simulations illustrate how cell-cell communication can be established via mechanical cues. Specifically, following active cell contraction, ECM realignment and deformation propagate mechanical cues. Firstly, Fig 6A (right panel) shows that cellular crosstalk is established via the formation of a continuous (percolative) path of maximal stress between both cells (see section 2.3). Furthermore, the stress distribution within the ECM shows higher stress levels aligned along the axis connecting the two cells.

**Fig 6.**
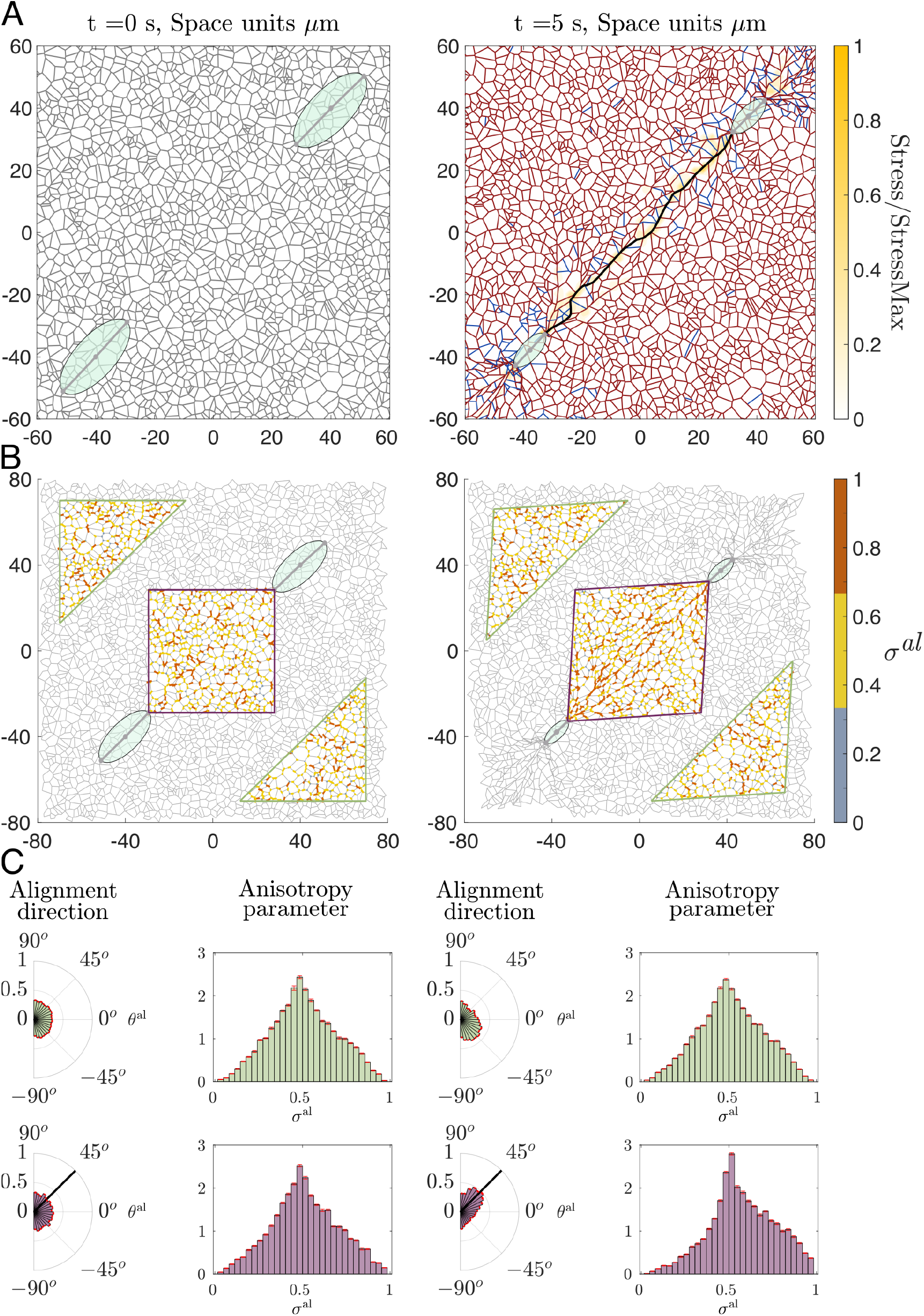
Contraction of two elongated cells. **(A)** Left panel: initial configuration of our model simulations in which two elongated cells are placed on a relaxed ECM. Right panel: the configuration of the cell-ECM system after *t* = 5 s of contraction. ECM fibres at equilibrium are shown in grey, extended fibres highlighted in dark red and compressed fibres in blue. The colour map represents the stress within the ECM, normalised by the maximum stress value. In addition, the ECM path of maximum stress transmission is highlighted in black. Parameters: Young’s modulus of the ECM fibres *E*_t_ = 100MPa, number of integrins per FA *N* = 1500, *F* ^co^ = 50nN. The remaining parameters are as listed in S1 Table. **(B)** Two plots comparing the local alignment of ECM fibres before and after cell contraction. For each crosslink, we calculate the principle alignment direction and the corresponding value of the anisotropy metric given by Eq (8). Orange color represents strong fibre alignment, yellow represents an intermediate anisotropy and grey indicates isotropic cases. We then visualise fibre alignment at crosslinks in a high interaction region between the two cells (purple rhombus), and in a low cell-cell interaction region (green rhombus). **(C)** Distributions of alignment direction and alignment measure in the high and low interaction regions outlined in **(B)**. Red error bars indicate the standard deviation calculated over 11 realizations of our model for the scenario of two contracting cells. Left panels show distributions of these metrics at equilibrium (*t* = 0 s), while right panels correspond to *t* = 5 s after contraction. Black solid lines in distributions of alignment angle indicate the direction of the segment that connects the cell centres (in purple region of high cell interaction). The t-test of the distribution of the anisotropy parameter in the region of high interaction before and after contraction indicates that the distributions are different (p-value = 1.9124 × 10^−04^). On the other hand, the t-test for the low interaction region before and after the contraction indicate that both samples could come from the same distribution with p-value = 0.7293.

Another crucial aspect that influences cell behaviour is the alignment of the ECM fibres, as cells tend to polarise along the matrix fibres and their migration direction can be biased by the local orientation of the ECM [56]. Our simulations, Fig 6B, indicate that while the realignment of the ECM following active cell contraction is heterogeneous, it is significantly stronger in the region of the ECM located between the active cells (outlined in purple) than in regions further apart from them (shown in green).

Using the quantifications of local alignment and local anisotropy defined in section 2.2, Eq (8), we quantify the ECM realignment in response to cell contraction. Fig 6C shows the distribution of metrics for ECM alignment anisotropy within the area between (far from) the contracting cells, represented by the purple rhombus (green triangles) in Fig 6B. Since the contraction-induced stress decays with distance, there is no significant change in the level of ECM anisotropy in the region of low cell interactions. By contrast, ECM fibres reorient towards the path of maximal stress that connects the two contracting cells. In the high interaction region (in purple), our model simulations predict the alignment of fibres in the direction of highest stress transmission (corresponding to 45^*°*^ in our simulation setup). Furthermore, the distribution of the local anisotropy, *σ*^*al*^, skews towards 1, which indicates an increase in ECM anisotropy in this region.

### 3.2 Parameter sensitivity analysis

#### Active contraction forces increase the force and stress transmitted between cells, but do not affect the kinetics of force transmission

A key parameter that determines the qualitative behaviour of the system is the active contractile force. Regulation of active contraction, specifically myosin motor activity, is crucial for controlling cell-substrate adhesion, cell migration and tissue architecture [8, 9, 36, 51, 57]. In order to analyse the role of the contraction force in the deformation of the substrate, we perform simulations in which we increase the cell contraction force within a range that maintains cell-ECM attachment. Specifically, the contraction force varies from *F* ^co^ = 10^4^ pN to *F* ^co^ = 5 · 10^4^ pN. The results of this sensitivity analysis are shown in Fig 7.

**Fig 7.**
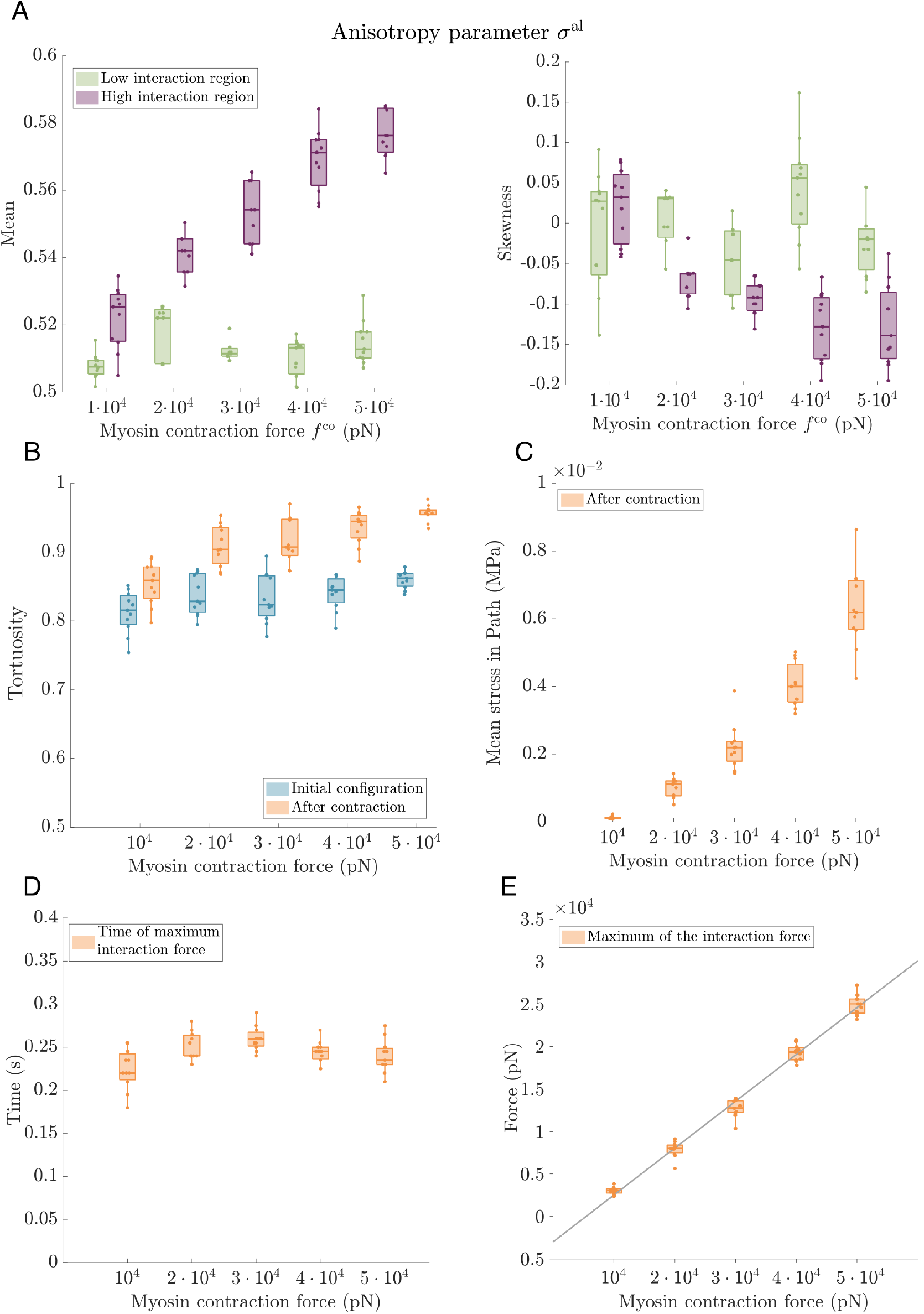
Parameter sweep analysis. Active contraction. We analyse the influence of active contractile forces on the mechanical communication between cells by measuring the alignment of ECM fibres, stress transmitted between cells and the peak of the interaction force at *t* = 5 s after contraction. In these simulations, the contraction force varies from *F* ^co^ = 10^4^ pN to *F* ^co^ = 5 · 10^4^ pN. Parameters: number of integrins per FA *N* = 1500, *E*_*t*_ = 10^7^ Pa. The remaining parameters are as listed in S1 Table. All results are averaged over 11 realisations of our model with random initial ECM network topologies. **(A)** Two boxplots showing the mean and skewness of the distribution of the anisotropy parameter for ECM fibres at *t* = 5 s after cell contraction. Green corresponds to the low interaction region while the results for the high interaction region are shown in purple (see Fig 6). **(B)** A plot showing the tortuosity of the path of maximum stress transmission (see section 2.3) calculated as a ratio of the Euclidean distance between the cells and the length of the percolation path. Orange indicates statistics calculated at *t* = 5 s after contraction, while the tortuosity of the same path at the initial time is shown in blue. **(C)** A plot showing the mean tensional stress in the maximum stress transmission path. **(D), (E)** Two plots demonstrating the time at which the cell-ECM interaction force,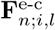, is maximum **(D)** and the value of this peak force **(E)** (see peak force in Fig 4C). In addition, we include a linear regression fit where we can see that the maximum of the interaction force grows linearly with the increment of the contractile forces. The fit has R-square = 0.9833.

Fig 7A-C show that the ECM becomes more aligned as the contractile forces increase. The behaviours of the mean (left panel) and skewness (right panel) of the local anisotropy parameter, Eq (8), are presented in Fig 7A, which shows that increasing the contraction force increases the anisotropy of the ECM mesh. In the region between the cells, elevating the contraction force increases both the mean anisotropy and the skewness of its distribution. Furthermore, in agreement with the fibre alignment results shown in Fig 7A, the path of maximal stress transmits more stress and becomes less tortuous as the contraction force increases.

Crucially, larger contraction forces enhance cell-to-cell communication. Fig 7E shows that the force exerted on the FAs increases as the contractility grows, augmenting the local deformation of the ECM. This local deformation is then propagated through the ECM, stretching and realigning the fibres between the cells, increasing the anisotropy of the fibres in the interaction region (purple, see Fig 6). While increasing the contractility enhances the transmission of forces through the ECM, the kinetics of the cell-to-cell communication are weakly affected. Fig 7D shows that the time at which the interaction force reaches its maximum, and its value (see Fig 6), are independent of the active contraction force.

#### ECM stiffness is crucial for determining the timing of mechanical communication between the cells

The effects of the mechanical properties of the ECM on cell behaviour are widely reported in the literature (see [3] for a review). In particular, ECM viscoelasticity plays an important role in cell migration and communication [20, 31, 37, 58]. Here we investigate how varying the stiffness of the ECM fibres affects mechanical communication between cells. Specifically, we conduct model simulations involving two contracting elongated cells in which we vary the Young’s modulus of the ECM fibres. In Fig 8, we present simulation results comparing the fibre alignment and stress transmission after active contraction of the two elongated cells.

**Fig 8.**
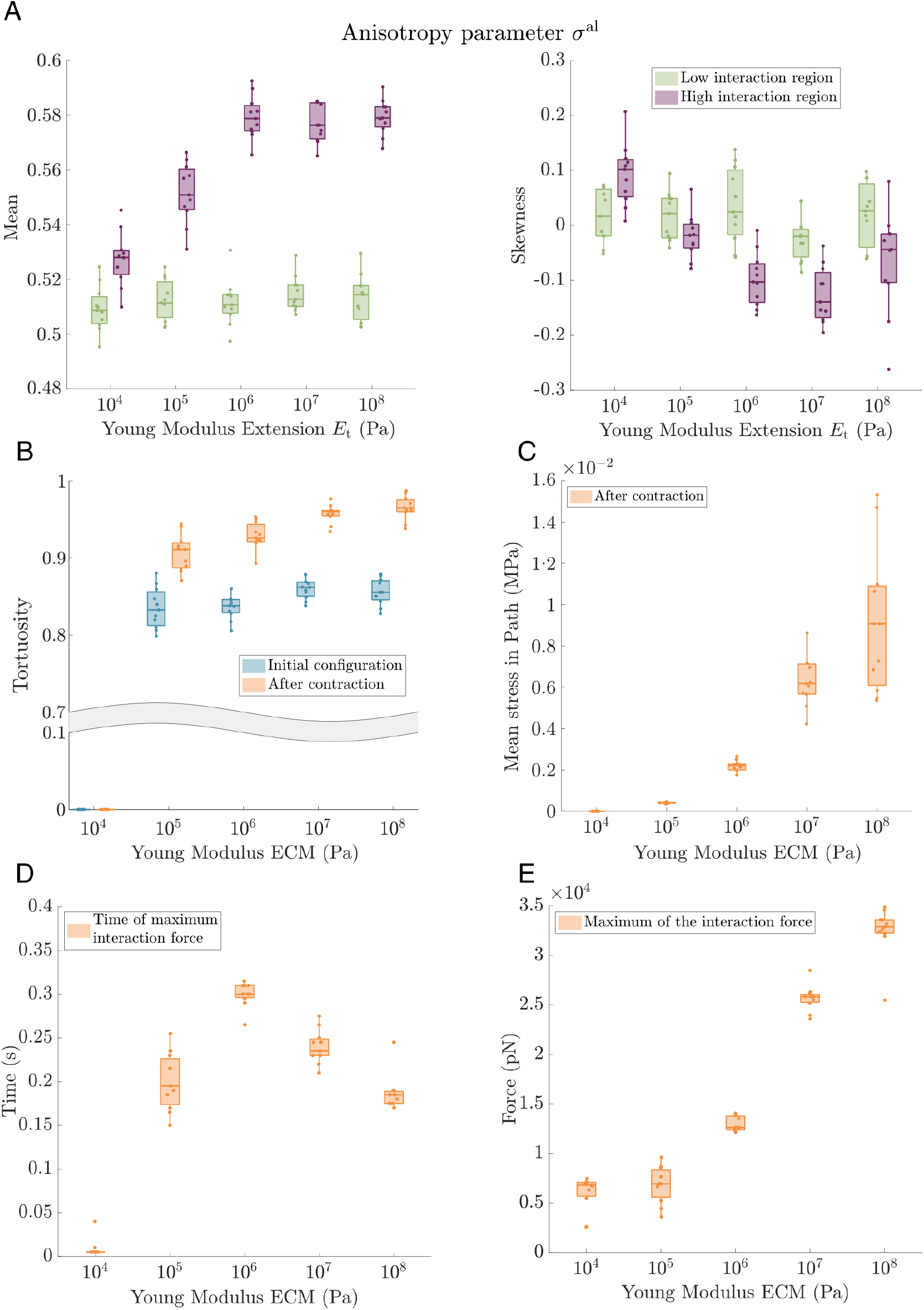
Parameter sweep analysis: Varying ECM stiffness. We analyse the influence of the stiffness of the ECM fibres on the mechanical communication between cells by measuring the alignment of the fibres, and the stress transmitted between cells at *t* = 5 s after cell contraction. ECM stiffness is varied from *E*_*t*_ = 10^4^ Pa to *E*_*t*_ = 10^8^ Pa. Parameters: number of integrins per FA *N* = 1500, *F* ^co^ = 50000 pN. The remaining parameters are listed in S1 Table. We perform 11 realizations with arbitrary initial configurations of the ECM for each set of parameter values. **(A)** We present the mean and skewness of the anisotropy parameter distribution of the ECM configuration at *t* = 5 s after cell contraction. Similar to results shown in Fig 6, green (purple) colour indicates the ECM region with low (high) cell interaction. **(B)** A boxplot showing the tortuosity of the path of maximum stress transmission (see section 2.3), calculated as a ratio of the Euclidean distance between the cells and the length of the percolation path. Orange indicates statistics calculated at *t* = 5 s after contraction, while the tortuosity of the same path at the initial time is shown in blue. For *E*_*t*_ = 10^4^ Pa, we do not observe the formation of a path between cells along stretched fibres. **(C)** A plot showing the mean tensional stress along the maximum stress transmission path at *t* = 5 s after cell contraction. Since no path of stretched fibres forms between the cells for the softest ECM, *E*_*t*_ = 10^4^ Pa, the mean stress in these cases is 0 Pa. **(D)** Time at which the interaction force, 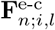, is maximum. **(E)** Maximum value of the interaction force. **(D)** and **(E)** show the time and the force value of the peak force represented in Fig 4C. In this case, the mean peak force is calculated from the mean of the interaction force of all the adhesion sites of both cells.

Fig 8A shows that anisotropy of the ECM mesh (both mean (left panel) and skewness (right panel)) within the high interaction region between cells increases with the Young’s modulus of its fibres, i.e. fibre realignment is transmitted better for the stiffer fibres. The ECM deformation saturates as the stiffness increases.

In Fig 8B-C we compare the statistics associated with the percolation path of maximum stress transmission described in section 2.3 as the Young’s modulus of the ECM fibres varies. For cases when a communication path along stretched fibres is formed, we calculate its tortuosity (Fig 8B) and the mean stress along its length (Fig 8C). Fig 8C shows that, for softer ECM, the mechanical deformation caused by cell contraction is not transmitted sufficiently far through the fibres to connect two cells separated 80 *µm* apart. Although the ECM fibres in the vicinity of the cell adhesion are stretched, the elastic forces resulting from this deformation are small, leading to less effective transmission of mechanical signals. Softer ECM also leads to less fibre realignment, with negligible reorientation after cell contraction for *E*_*t*_ = 10^4^ Pa. In the softest case, we performed a longer simulation, which showed that the mechanical percolation occurs at a slower rate, see S1 Fig in S1 File. On the other hand, fibre realignment and mechanical stress are better transmitted for stiffer ECM. Despite the saturation in the mean and skewness of the anisotropy parameter for stiffer ECM fibres (see *E*_*t*_ = 10^7^ Pa and *E*_*t*_ = 10^8^ Pa in Fig 8A), a higher Young’s modulus results in higher deformation and greater mean stress along the percolation path, see Fig 8B-C. For stiffer ECM, when contracting cells pull on ECM crosslinks, as the fibres are more rigid, small fibre stretch implies transmission of larger forces within the network.

Finally, in order to quantify the dynamics of the interaction force, in Fig 8D,E we determine the time at which the force resulting from the integrin stretching is maximum, and the value of this force (maximum of the interaction force illustrated in Fig 4C). For stiffer ECM fibres, increasing values of the Young’s modulus reduce the time at which the peak force appears and increase the peak force. Small deformations of the ECM fibres lead to tensile stresses (see in Figs 8C and E), and more rapid transmission of deformation through the ECM (see Fig 8D).

### 3.3 Local ECM topology influences its effective stiffness

For larger values of the Young’s modulus, cells frequently detach from the ECM during contraction. As cells pull on stiffer substrates, the peak force experienced by the integrin cluster is reached more rapidly, leaving insufficient time for the integrin binding to adjust. Consequently, cells enter the detachment basin of attraction (see Fig 5), even though they are experiencing lower forces than the critical force described in section 2.5. We posit that the time required to reach peak force at a FA is closely related to the topology of the ECM network. If the initial ECM configuration allows for the formation of a communication path, then cells remain attached because mechanical stress can be rapidly transmitted along this path, with the energy from the initial force dissipated through ECM deformation. On the other hand, if the initial ECM configuration lacks a clear communication path, then the network topology induces rigidity, preventing transmission of the deformation caused by the cell’s initial pull. This topological effect is illustrated in S2 Fig in S1 File, where we present two model configurations for which a cell detaches from the ECM, and one model configuration for which the cells do not detach. In these cases, the stress is transmitted via several paths. In cases for which a cell detaches, the initial cell contraction stretches the integrins that form FAs rather than the ECM fibres, forcing the FA system to enter the basin of attraction of detachment (see Fig 5C, the detachment region).

To further investigate how the local structure of the ECM network affects the stress exerted on the FAs, we perform simulations in which two cells attached to a simplified ECM are pulled away from each other (see Fig 9). We consider two ECM topologies that connect the two cells: a zigzag ECM path (see Fig 9A) and a more complex scenario where the cells are connected to two long fibres with small transversal fibres between them (Fig 9B,C). We assume that cells are attached to the ECM fibres at one of their adhesion sites, while being pulled away from each other with a constant force. We assume further that at the initial configuration, there are a certain number of bound integrins at FA sites. We use this setup to investigate how the ECM topology affects cell-ECM interactions as the ECM stiffness varies.

**Fig 9.**
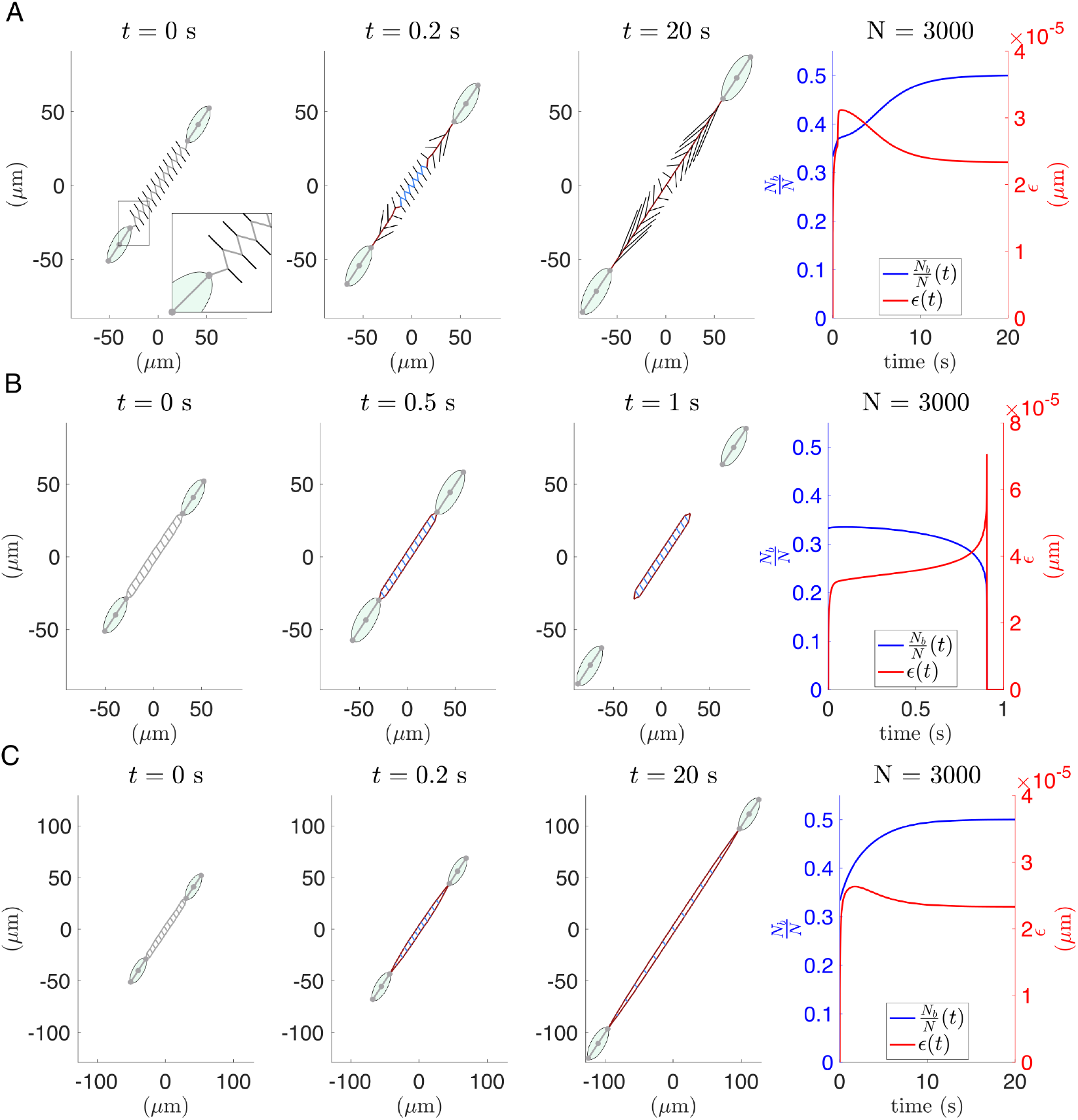
Two-cells simulation: analysis of topological effects of the ECM network. Numerical simulations with two cells connected to simplified ECM networks. We present three snapshots of the time evolution of the cell-ECM system, and the evolution of the key variables that define the interaction force: the proportion of bound integrins 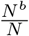 and their stretch *ϵ*^i^. The cells are pulled away from each other with a force *F* ^pu^ = 35 nN. In light grey, we plot the fibres that are in their equilibrium state, blue indicates compressed and red corresponds to stretched fibres. Rightmost panels show the dynamics of the proportion of bound integrins (blue) and their stretch (red). **(A)** Two cells connected by a single zigzag path of ECM fibres. The crosslinks of the main zigzag path are connected to lateral fibres with fixed outer endpoints. These fibres are plotted in black. In this case, the system achieves an equilibrium, where the ECM fibres are stretched, and they form a communication path between cells that remain attached to it. The Young’s modulus of the ECM fibres is *E*_*t*_ = 10^8^ Pa. **(B)** Simulation with the ECM configuration consisting of two paths connected by transversal fibres. Rigidity of this ECM configuration results in the inability of the system to form a communication path between cells. This leads to their detachment due to the rapid stretching of the integrins. The ECM fibres have the same Young’s modulus as in **(A)**, *E*_*t*_ = 10^8^ Pa. **(C)** The same ECM configuration as in **(B)** but with softer fibres, *E*_*t*_ = 10^6^ Pa. In this case, the cells do not detach from the ECM.

Fig 9A shows that, for the zig-zag scenario, the stress generated by the pulling force is dissipated through the deformation of the communication path, regardless of ECM stiffness. This deformation is rapidly transmitted, and the pulling energy dissipated by deforming the ECM. The FA system evolves until it reaches an equilibrium with a positive number of bound integrins (see right panel of Fig 9A). By contrast, the ECM configuration presented in Fig 9B introduces extra rigidity into the system, resulting in an entirely different behaviour. Since this ECM network is less deformable at the same ECM stiffness, its effective stiffness is greater and the pulling force leads to rapid stretching of the integrins that form the FAs, which causes cell detachment from the ECM (see Fig 9B, right panel). Additionally, in S1 Video, we observe that the FA system enters the detachment basin of attraction, even though the cell-ECM interaction force is below its critical value. There is no difference in behaviour when we randomly remove the transversal fibres (see S3 Fig in S1 File), which means that the topological effects result from the branching of the stress transmission in different paths.

Cell detachment due to the local structure of the ECM is closely linked to the effective stiffness of the ECM mesh. In Fig 9C, we present a simulation for a softer ECM where the cells remain anchored to the ECM. As the fibres are softer, the force exerted at FAs is transformed into substrate deformation, allowing part of the energy to be dissipated without over-stretching the integrins.

## 4 Discussion

In this work, we have presented an agent-based model that simulates mechanical interactions between cells and ECM fibres and captures the simultaneous time evolution of the ECM, cells and the FA. The model allows us to quantify cell-cell communication mediated by ECM deformation, and shows how this process depends on the mechanical properties of cells and ECM fibres and the topology of the ECM network.

Motivated by previous studies [14, 26–28], our analysis of the FA interaction system provides a detailed understanding of how binding-unbinding dynamics change in response to changes in externally applied forces. By coupling cell and ECM deformation with cell-ECM interactions, we use the model to simulate the temporal evolution of FA components during cell contraction.

The polygonal cell geometry used in our model enables us to study cells with a variety of shapes. For example, in Fig 10A, we observe the stress transmission path between two contracting fan-shaped cells. Our model can also accommodate several cells in a straightforward manner (see Fig 10B), enabling us to study the feasibility of indirect cell-cell communication mediated mechanical cues in the ECM.

**Fig 10.**
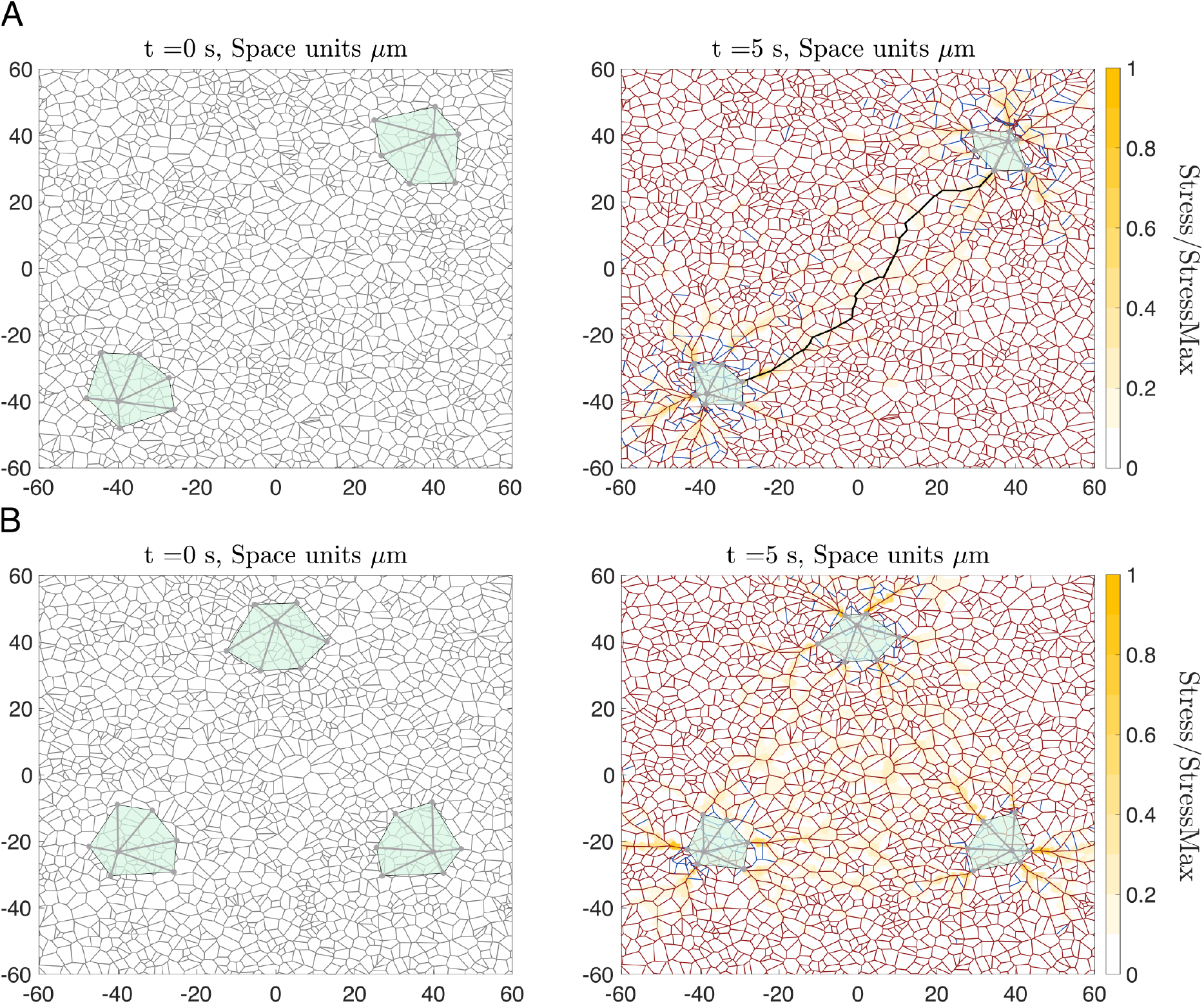
Illustration of the model’s flexibility to accommodate various cells and different cell shapes. Two model simulations with contracting fan-shaped cells. ECM fibres at equilibrium are shown in grey, extended fibres are highlighted in dark red and compressed fibres in blue. The underlying colour map indicates the stress within the ECM, normalised by the maximum observed stress. Young’s modulus of the ECM fibres was set to *E*_t_ = 1MPa, the number of integrins per FA *N* = 1000 and the contraction force *F* ^co^ = 6.6 nN. The remaining parameters are listed in S1 Table. **(A)** A numerical experiment similar to the one shown in Fig 6 performed for fan-shaped cells. Left panel: initial configuration in which two fan-shaped cells are placed on a relaxed ECM. Right panel: the configuration of the cell-ECM system after *t* = 5 s of contraction. The ECM path of maximum stress transmission between two cell interaction sites is highlighted in black. **(B)** Model simulation as in **(A)** but with three fan-shaped cells.

In this work, we used several metrics to describe the stress distribution and fibre alignment within the ECM. These metrics include: the anisotropy parameter, alignment direction and the stress of individual fibres. By computing these metrics we study how ECM stiffness and cell contraction forces affect cell-cell communication.

Many open questions remain regarding ECM deformation and the mechanical communication between cells [4, 21, 48]. Our model introduces novel aspects with respect to previous agent-based models [31–33, 37, 47] in two ways: (*i*) we introduce a novel force-based model that incorporates the internal dynamics of cell structure, cellular contractility and FA regulation, which mediates cell-ECM interaction, to an elastic network description for the ECM fibres; (*ii*) by integrating these processes—which range from subcellular activities (such as myosin-driven contraction and the mechanosensitive binding and unbinding of integrins) to ECM deformation—we can analyse and compare the different timescales associated with them and their impact on overall dynamics.

Several experimental studies have shown that cell behaviour is sensitive to the stiffness of the ECM [4, 15, 16, 30, 48, 59, 60]. Our results indicate that for soft substrates the rate of transmission of deformation within the ECM slows down, hindering the mechanical communication between cells (see Fig 8 and S1 Fig in S1 File). Furthermore, in agreement with previous experimental studies [4, 15, 48], our model predicts that increasing substrate stiffness significantly facilitates cellular mechanosensing as there is a faster transmission of the deformation, which is associated with a greater stress in the ECM (see Fig 8). In addition, our modelling results confirm experimental studies [30], which show that in stiff substrates the integrins unbind from the ECM, preventing the force transmission from the cells to the ECM (see Fig 9 and S2 Fig in S1 File). Furthermore, our model shows that the ECM configuration can generate an increased *effective* rigidity, i.e. for ECM fibres with the same Young’s modulus, the network topology affects its ability to deform under applied forces. Fig 9 and S2 Fig in S1 File show how the *effective* rigidity can causes cell detachment from stiff ECM, those with a Young’s modulus of O(100 MPa), i.e. in simulations involving fibres with the same Young’s modulus, the detachment depends on the ECM configuration. We conclude that, as for soft substrates, the transmission of deformation slows down, and cell detachment is more easily facilitated on stiff substrates; an ECM with intermediate stiffness will maximise mechanical communication between cells. Several experimental studies have investigated how active mechanical cues such as contractile forces [59] and intrinsic properties of the matrix, for example the stiffness of their fibres [60], affect matrix reorganisation. Our model describes how contractile forces and the ECM stiffness affect the transmission of cell-generated forces through the ECM (see Figs 7 and 8 respectively).

Our model integrates multiple biophysical processes which act on multiple temporal scales. We consider simultaneously the dynamics of sub-cellular structures (e.g cytoskeleton and myosin motors), the deformation of ECM fibres, and the regulation of FAs, which coordinate the transmission of bio-mechanical signals between the ECM and sub-cellular structures. The simulation results in Figs 4B-C predict that the binding-unbinding of integrins within a FA occurs on a slower timescale than that associated with their deformation. This indicates that the dynamics of cell-ECM interactions may be very sensitive to the stretching of integrins. The forces governing integrin deformation mainly arise from cellular contractility, cell resistance to deformation, and substrate elasticity, which act on timescales similar to those associated with the integrin dynamics. Therefore, the evolution of FA components and cell-ECM interactions is largely determined by how integrins respond to these forces. Our model simulations suggest that cell detachment from a stiff ECM occurs because the mechanical properties of the ECM limit its deformation, causing integrins to stretch too quickly. In turn, rapid integrin deformation causes the integrins to unbind from the ECM, leading eventually to cell detachment (see Fig 9 and S1 Video). The simulations shown in S2 Fig in S1 File, S2 Video, and S4 Video demonstrate that cell detachment from the ECM can occur even when the forces acting on the FAs are below the critical value if the force due to cell contraction drives the FA system to enter the detachment basin of attraction (Fig 5C). In contrast, when cells do not detach, the FA system evolves to a stable equilibrium, with a positive number of bound integrins (S3 Video). In conclusion, our model shows that non-steady-state effects, where forces evolve due to the dynamic interplay between the cells and the ECM, play a crucial role in regulating cell-ECM interactions. Thus, to accurately capture the mechanical details of cell-ECM interactions, it is essential to consider the simultaneous evolution of these different components.

Although our model captures the detachment of FAs and ECM deformation, more research is needed to better describe the cell-ECM crosstalk and its influence on cell behaviour. A natural progression of this work is to incorporate related cellular and ECM processes, including ECM remodelling, which modify the mechanical properties of the ECM [33], and the de novo formation of cell-ECM bonds following cell detachment from the ECM, an essential model extension for simulating cell migration [40].

In summary, our model represents the first step in the development of a mechanical framework for modelling cell-ECM interactions. This framework aims to integrate processes acting on different spatial and temporal scales to understand what regulates cell-cell communication and how mechanical feedback influences cell behaviour.

## Acknowledgments

This work has been funded by the Spanish Research Agency (AEI), through the Severo Ochoa and María de Maeztu Program for Centers and Units of Excellence in R&D (CEX2020-001084-M). D.S. and T.A. thank CERCA Program/Generalitat de Catalunya for institutional support. J.A-T, D.S. and T.A. have been funded by grant PID2021-127896OB-I00 from MCIN/AEI/10.13039/501100011033 ‘ERDF A way of making Europe’. J.A-T is supported by a scholarship from the program ‘Ayudas para Contratos Predoctorales para la Formaciín de Doctores’ (PRE2021-099696).

