## Supplementary material for "Multiscale modelling shows how cell-ECM interactions impact ECM fibre alignment and cell detachment": S1 File

**S.3 Full system of model equations and parameter values** **9**

**S.4 Numerical methods** **10**

**S.5 Exact calculation of the critical value $F^{\mathbf{BP}}$** **11**

### S.1 Supplementary Figures

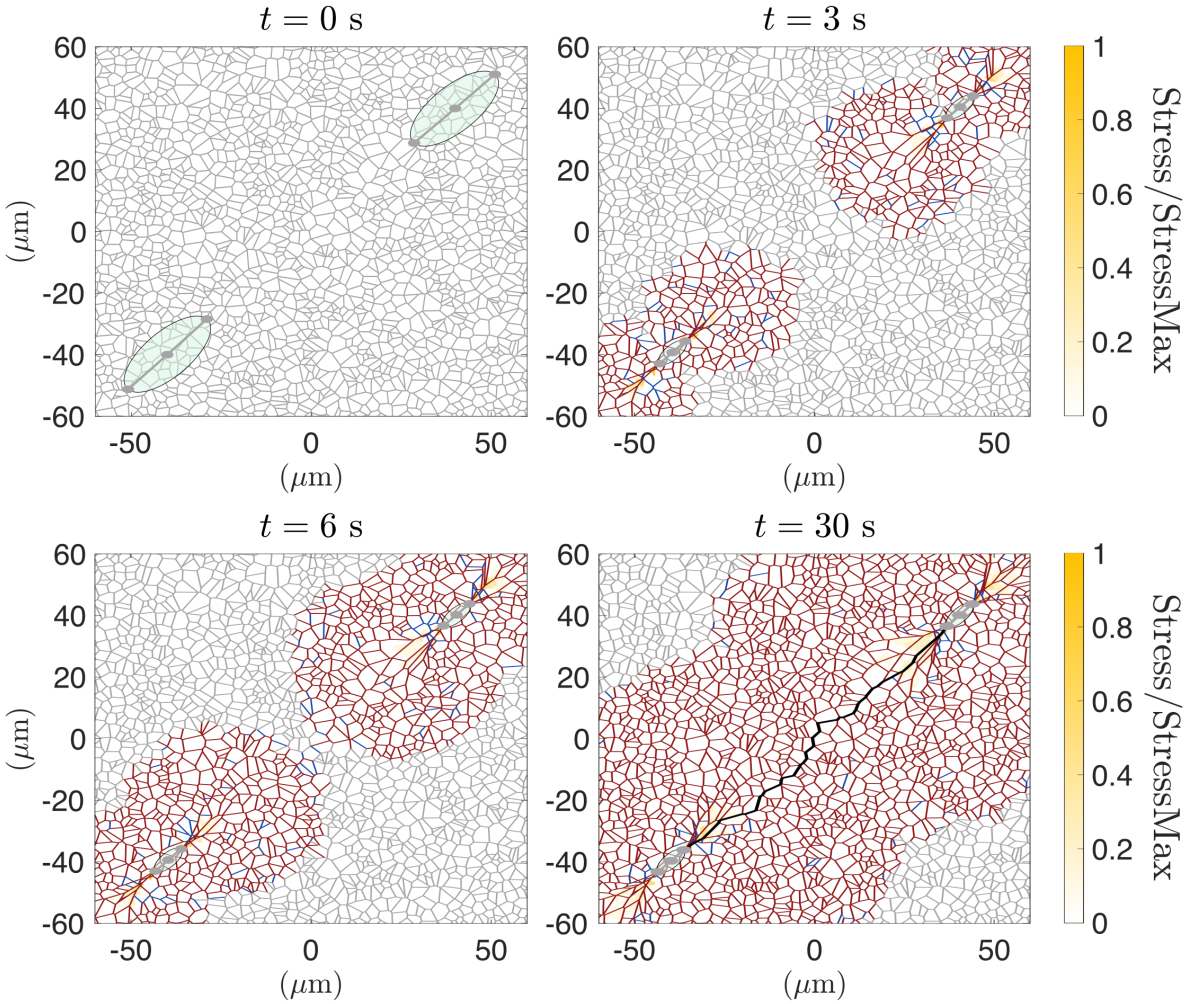

**Fig S1. Force transmission on soft ECM. Longer time simulation.** Simulation with two cells contracting until $t = 30s$ on a soft ECM, i.e. $E_t = 10^4$ Pa. ECM fibres at equilibrium are shown in grey, extended fibres highlighted in dark red and compressed fibres in blue. The colour map represents the stress within the ECM, normalised by the maximum observed stress.

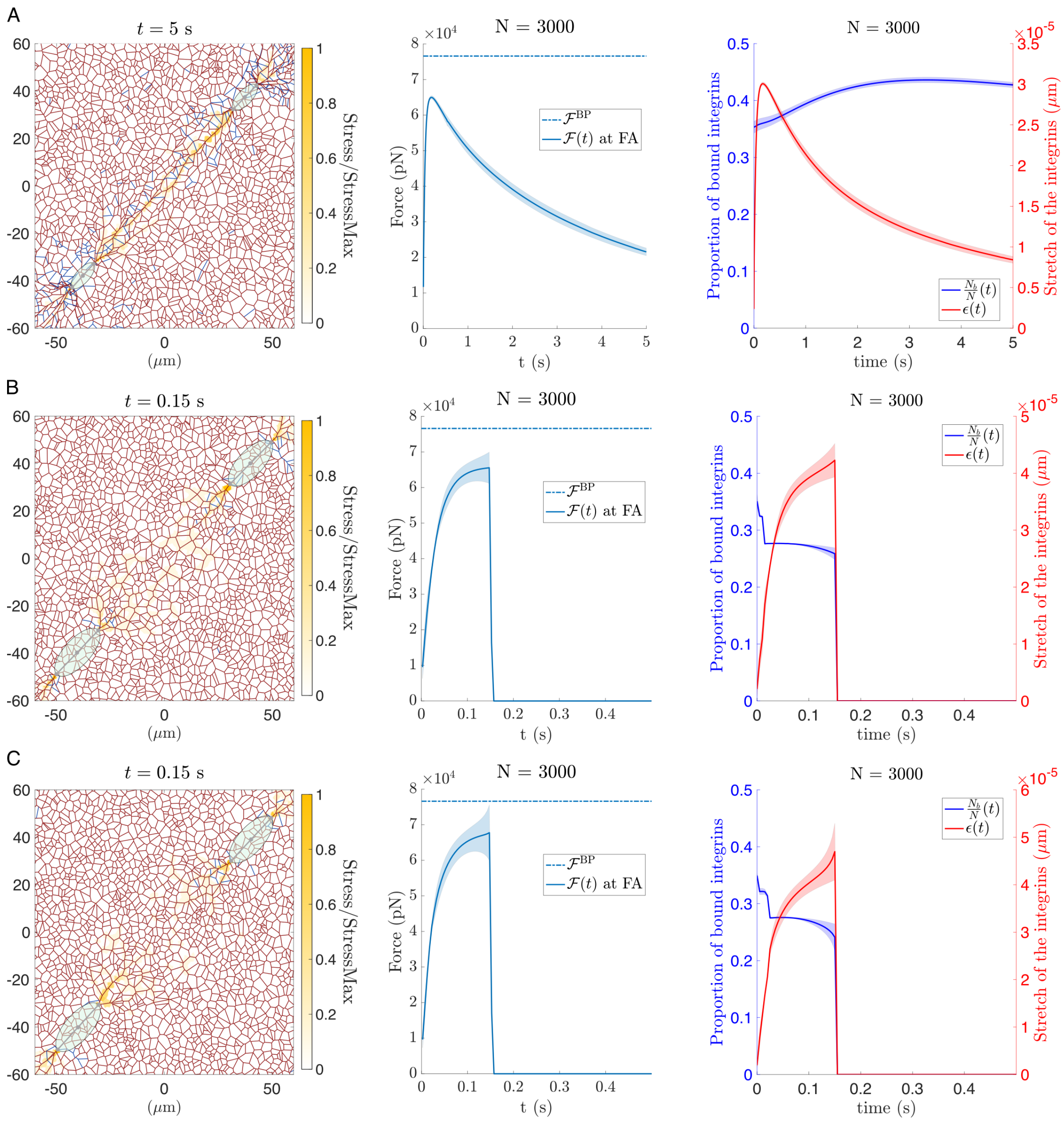

**Fig S2. Force generation on stiff ECM.** Three simulations where cells contract on a stiff ECM, i.e. $E_t = 10^8$ Pa. Left panel: configuration corresponding to the moment before detachment. ECM fibres at equilibrium are shown in grey, extended fibres highlighted in dark red and compressed fibres in blue. The colour map represents the stress within the ECM, normalised by the maximum observed stress. Middle panel: we include the evolution of the force exerted on the FA and the value of the critical force (details in 2.5 of the main text). Right panel: we include the evolution of the proportion of bound integrins in blue and their stretch on red. We average over the FA facing both cells. **(A) Test 1** Two cells contracting, similar to the scenario presented in Fig 6. **(B) Test 2** We can see in yellow the stress transmission paths between the cells. In the top right cell, we see how two paths emerge, which causes the extra rigidity in the ECM. **(C) Test 3** In this configuration, the two stressed path emerge in the bottom left cell.

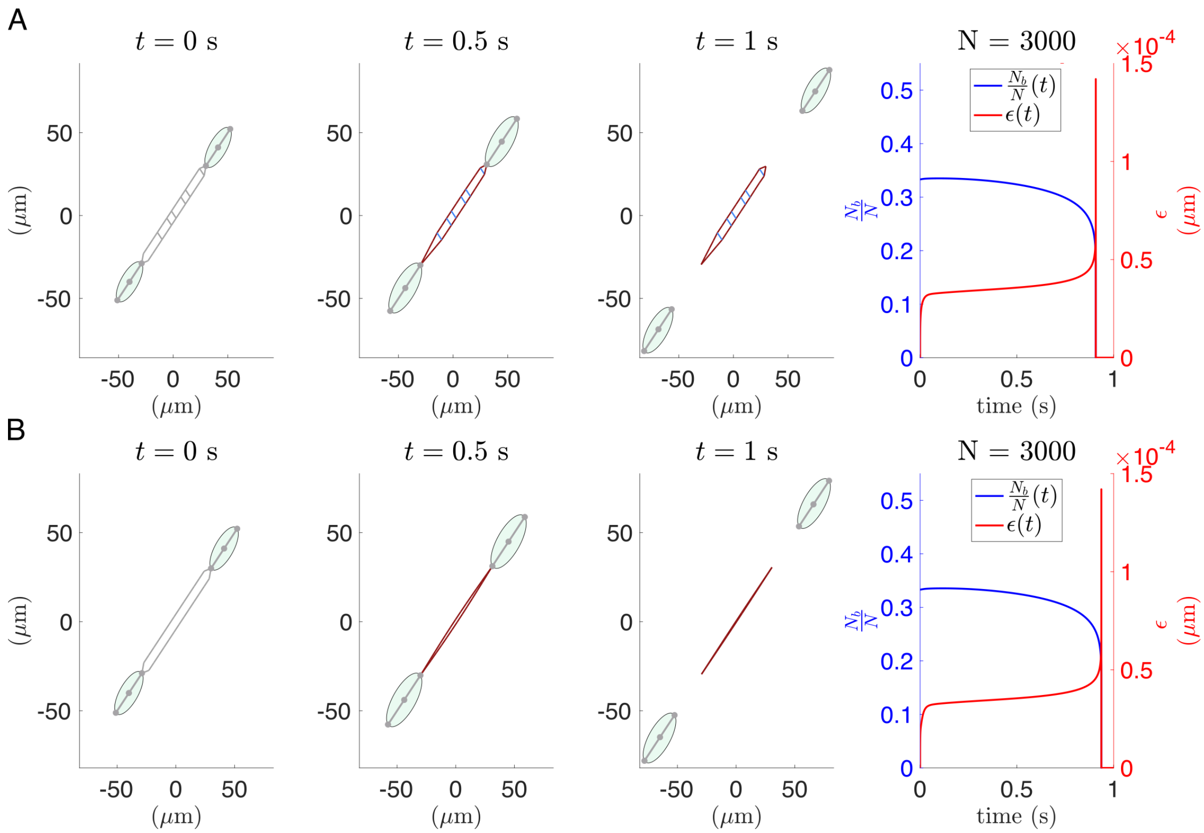

**Fig S3. Random removal of transversal fibres. Detachment of stiff substrate** Simulation with two cells that detach from a stiff substrate, i.e. $E_t = 10^8$ Pa. Similar configuration as in Fig 9B, but including the random removal of transversal fibres. The qualitative behaviour of the system in this case does not change, the cells detach from the ECM.

### S.2 Detailed model description

Our model consists of three main parts describing the dynamics of the ECM, cells, and their interaction. Here, we present a more detailed description of the model's building blocks.

#### S.2.1 Modelling the deformation of the ECM as a mesh of elastic fibres

Following [1], we consider a simple mechanical model to describe the dynamics of the substrate. We model the ECM as a network whose edges represent individual ECM fibres, which are assumed to behave as linear springs (see Fig 1A). Individual fibres interconnect through the formation of molecular bonds called crosslinks. The distribution of crosslinks confers the overall connectivity and complex rheological properties of the ECM [1–3].

Our model tracks the positions of the set of crosslinks, $\Omega_{\mathrm{e}}$, by solving a set of ordinary differential equations (ODEs). Each crosslink is connected to a set of neighbouring nodes (crosslinks), $\Omega_{\mathrm{e},l}$. The evolution of the ECM nodes depends on the elastic forces acting upon them due to the deformation of individual ECM fibres (Fig 2A left panel) as well as the cell-ECM interaction forces at the adhesion sites (Fig 2B). In the overdamped limit, the position of the $l$-th ECM crosslink, $\mathbf{x}_i^{\mathrm{e}}$, of the network is given by:

$$\eta \frac{d\mathbf{x}_l^{\mathrm{e}}}{dt} = \mathbf{F}_l^{\mathrm{el}} + \mathbf{F}_{l,n;i}^{\text{e-c}}. \tag{1}$$

Here, the elastic force resulting from the deformation of individual fibres connected to the $l$-th crosslink, $\mathbf{F}_l^{\mathrm{el}}$, depends on the local structure of the ECM (i.e. neighbouring nodes) and its elastic properties:

$$\mathbf{F}_l^{\mathrm{el}} = \sum_{m\in\Omega_{\mathrm{e},l}} -k_{l,m}^{\mathrm{el}}(\|\mathbf{x}_l^{\mathrm{e}} - \mathbf{x}_m^{\mathrm{e}}\| - L_{l,m}^0)\frac{\mathbf{x}_l^{\mathrm{e}} - \mathbf{x}_m^{\mathrm{e}}}{\|\mathbf{x}_l^{\mathrm{e}} - \mathbf{x}_m^{\mathrm{e}}\|}. \tag{2}$$

where $L_{l,m}^0$ denotes the natural length of the fibre connecting nodes $l$ and $m$ ( with positions $\mathbf{x}_l^{\mathrm{e}}$ and $\mathbf{x}_m^{\mathrm{e}}$ respectively) and $k_{l,m}$ is the corresponding elastic constant. The elastic constant $k_{l,m}^{\mathrm{el}}$ of each fibre is determined by its corresponding Young's modulus, $E_{l,m}$, cross-sectional area, $A_{l,m}$, and natural length $L_{l,m}^0$:

$$k_{l,m}^{\mathrm{el}} = \frac{E_{l,m}A_{l,m}}{L_{l,m}^0}. \tag{3}$$

For simplicity, we assume that all ECM fibres have the same Young's modulus, $E$, and constant cross-sectional area, $A$. Nonetheless, our model can easily be extended to account for the heterogeneity and diferent shape properties of individual matrix fibres. Following [1], we account for the softer compressive response of the fibres by assigning two values to the Young's modulus depending on extension or compression:

$$E = \begin{cases} E_{\mathrm{t}}, & \text{fibre extension,} \\ E_{\mathrm{c}}, & \text{fibre compression,} \end{cases}.$$

with $E_\mathrm{t} > E_\mathrm{c}$.

#### S.2.2 Modelling cell shape and its deformation

Cell shape must be robust to random perturbations but also adaptive to facilitate vital cellular functions, such as migration and interaction with the external environment. The cell cytoskeleton plays a major role in organising the cellular content and providing structural support, enabling cells to adapt their shape according to their state and function [4]. The cytoskeleton is formed by three main types of polymers: actin filaments, microtubules and intermediate filaments, and regulatory proteins. These polymers form complex networks that can resist deformation and reorganize in response to applied forces [4]. Viscoelastic properties at the cell level emerge from the elastic properties of the cytoskeleton polymers and their connectivity [5–7].

Rather than addressing the full complexity of the cytoskeleton dynamics and cell shape, we propose a simple model that captures the main mechanical properties of cells. In particular, cells are represented as collections of viscoelastic elements that connect the cell's organising centre with the sites of FA assembly. These viscoelastic elements can be identified with actin bundles or stress fibres. Fig 1A illustrates different cell geometries that can be obtained by varying the number and the length of these viscoelastic elements. As Fig 1A shows, this representation can account for cell polarity: rounded vs elongated cell shape, as well as front-to-back symmetry breaking. Using this framework, we can describe an arbitrary number of cells $N_\mathrm{c}$. The shape of the $n$-th cell, $n \in 1, \ldots, N_\mathrm{c}$, is then determined by the positions of the set of points: $\left(\mathbf{x}^\mathrm{c}_{n;i}\right)_{i=0}^{N^\mathrm{c}_n}$, $\mathbf{x}^\mathrm{c}_{n;i} \in \mathrm{R}^2$, with $\mathbf{x}^\mathrm{c}_{n;0}$ representing the position of the cytoskeleton organising centre, and $\mathbf{x}^\mathrm{c}_{n;i},\ i \in \Omega^\mathrm{c}_n = 1, \ldots, N^\mathrm{c}_n$ describing the positions of the end points of the stress fibres, where the FAs assemble. $N^\mathrm{c}_n$ represents the number of adhesion sites in the $n$-th cell.

Fig 2C illustrates the forces that account for cellular geometry: structural forces, active forces generated by myosin motors, and forces exerted by the ECM. In the overdamped limit, where viscous effects dominate over inertial ones, $\left(\mathbf{x}^\mathrm{c}_{n;i}\right)_{i=0}^{N^\mathrm{c}_n}$ are determined by the balance of all the forces acting upon the system. The resulting set of ODEs describing the evolution of $\left(\mathbf{x}^\mathrm{c}_{n;i}\right)_{i=0}^{N^\mathrm{c}_n}$ are:

$$
\begin{aligned}
\eta \frac{d\mathbf{x}^\mathrm{c}_{n;0}}{dt} &= -\sum_{i \in \Omega^\mathrm{c}_n} \left(\mathbf{F}^\mathrm{st}_{n;i} + \mathbf{F}^\mathrm{co}_{n;i}\right), \\
\eta \frac{d\mathbf{x}^\mathrm{c}_{n;i}}{dt} &= \mathbf{F}^\mathrm{st}_{n;i} + \mathbf{F}^\mathrm{co}_{n;i} + \mathbf{F}^\mathrm{e\text{-}c}_{n;i,l} \qquad \text{for } i \in \Omega_c.
\end{aligned}
\tag{4}
$$

The structural forces $\mathbf{F}^\mathrm{st}_{n;i}$ correspond to the physical support provided by the cell cytoskeleton. In this work, we consider two mechanical contributions of the cytoskeleton, one associated with the structural support provided by the stress fibres, $\mathbf{F}^\mathrm{kv}_{n;i}$, and the other, the so-called *angular forces*, $\mathbf{F}^\mathrm{an}_{n;i}$, which aim at maintaining cell shape. These angular forces act to maintain the target or natural angle between neighbouring stress fibres, similarly to elastic forces that aim to maintain the natural length of elastic springs:

$$
\mathbf{F}^\mathrm{st}_{n;i} = \mathbf{F}^\mathrm{kv}_{n;i} + \mathbf{F}^\mathrm{an}_{n;i}. \tag{5}
$$

Regarding the former, we assume that stress fibres are viscoelastic materials, and, following previous work [8], we assume that their deformation can be described by the Kelvin-Voigt model. This model describes the behaviour of a system equivalent to a linearly elastic element in parallel with a linear viscous dashpot. Accordingly, the stress (force) exerted by the stress fibre on its end point is given by:

$$\begin{aligned}
\mathbf{F}^{\mathrm{kv}}_{n;i} &= \left(-\kappa^{\mathrm{el}}_i \varepsilon^{\mathrm{c}}_{n;i} - \kappa^{\mathrm{vi}}_i \frac{d\varepsilon^{\mathrm{c}}_{n;i}}{dt}\right)\hat{\mathbf{x}}_{n;i} \\
\varepsilon^{\mathrm{c}}_{n;i} &= L^{\mathrm{c}}_{n;i} - l^{\mathrm{c}}_{n;i}, \\
L^{\mathrm{c}}_{n;i} &= \left\|\mathbf{x}^{\mathrm{c}}_{n;i} - \mathbf{x}^{\mathrm{c}}_{n;0}\right\|, \\
\hat{\mathbf{x}}_{n;i} &= \frac{\mathbf{x}^{\mathrm{c}}_{n;i} - \mathbf{x}^{\mathrm{c}}_{n;0}}{\left\|\mathbf{x}^{\mathrm{c}}_{n;i} - \mathbf{x}^{\mathrm{c}}_{n;0}\right\|},
\end{aligned} \tag{6}$$

where $l^{\mathrm{c}}_{n;i}$ is the natural length, $\kappa^{\mathrm{el}}_i$ is the elastic constant and $\kappa^{\mathrm{vi}}_i$ is the damping constant of the $i$-th viscoelastic element of the cell.

The angular force is derived from a potential energy that establishes a preferred angle between two consecutive stress fibres and penalises deviations from it. The resulting restoring force is given as follows:

$$\begin{aligned}
\mathbf{F}^{\mathrm{an}}_{n;i} &= -\nabla_i \left(\sum_{j\neq i, j\in\Omega^{\mathrm{c}}_n} \frac{1}{2}\kappa^{\mathrm{be}}_{n;i,j}(\cos\theta_{n;i,j} - \cos\theta^0_{n;i,j})\right), \\
\cos\theta_{n;i,j} &= \langle \hat{\mathbf{x}}_{n;i}, \hat{\mathbf{x}}_{n;j}\rangle,
\end{aligned} \tag{7}$$

where $\theta^0_{n;i,j}$ is the natural angle between the segments $(\mathbf{x}^{\mathrm{c}}_{n;0} - \mathbf{x}^{\mathrm{c}}_{n;i})$ and $(\mathbf{x}^{\mathrm{c}}_{n;0} - \mathbf{x}^{\mathrm{c}}_{n;j})$.

Active contraction forces play a crucial role in cellular dynamics and intracellular cytoskeleton organisation [9]. The main source of cortical contractibility at the cellular level are the non-muscle myosin II motors [7, 10], which act at the actin filament networks. In our model, we account for the effects of myosin contraction by incorporating a constant contraction force acting on the cell's structural segments:

$$\mathbf{F}^{\mathrm{co}}_i = F^{\mathrm{co}}\hat{\mathbf{x}}_{n;i}. \tag{8}$$

#### S.2.3 Focal adhesion dynamics: cell-ECM interaction forces

Focal adhesions are protein complexes that mediate the interaction between cells and the ECM. One of the main components of these adhesion structures are integrins [11, 12], transmembrane receptors that link cytoskeleton actin bundles with the ECM [13]. Integrins mediate the transmission of mechanical forces due to cell-ECM interaction [12]. The formation of integrin clusters at FA sites is regulated internally by the cell and depends on several factors, including the organisation of actin bundles, cellular contraction forces, and the force exerted on the cell by the ECM [13]. Our model considers a fixed number of integrins per FA, which can be either bound to one of the ECM crosslinks or disconnected from the ECM (unbound). We represent the mechanical properties of a cluster of bound integrins as a set of linear springs connected in parallel, which allows us to assume that the intensity of the cell-ECM interaction force is directly proportional to the number of bound integrins. The formation of bonds between the ECM and unbound integrins is assumed to occur at a constant rate, whereas the unbinding rate is force-dependent [14, 15].

We denote by $N^b_{n;i,l}(t)$ the number of bound receptors forming links between cell vertex $i$ and crosslink $l$ of the cell $n$. We further assume that the total number of integrins within the FA at vertex $i$, $N_{n;i}$, is conserved and that each bound integrin behaves as a Hookean spring. Since we assume that integrins form an in-parallel array, all the bound integrins within the same FA experience the same stretch, $\epsilon^{\mathrm{i}}_{n;i,l}(t)$, which is determined by the distance between the ECM crosslink and the cell adhesion site:

$$\epsilon^{\mathrm{i}}_{n;i,l}(t) = \left( \left\| \mathbf{x}^{\mathrm{c}}_{n;i} - \mathbf{x}^{\mathrm{e}}_{l} \right\| - L^{\mathrm{i}}_{0} \right).$$

Thus, the total force exerted by a cluster of bound integrins at $\mathbf{x}^{\mathrm{c}}_{n;i}$ is given as follows:

$$\mathbf{F}^{\text{e-c}}_{n;i,l} = -N^b_{n;i,l} k^{\mathrm{i}} \epsilon^{\mathrm{i}}_{n;i,l} \frac{\mathbf{x}^{\mathrm{c}}_{n;i} - \mathbf{x}^{\mathrm{e}}_{l}}{\left\| \mathbf{x}^{\mathrm{c}}_{n;i} - \mathbf{x}^{\mathrm{e}}_{l} \right\|}. \tag{9}$$

The evolution of $N^b_{n;i,l}$ is prescribed by a continuous binding-unbinding process of integrins where the unbinding process is assumed to be mechanosensitive, i.e. the unbinding rate is a function of the total force exerted on the integrins [16, 17]:

$$F^{\mathrm{i}}_{n;i,l} = k^{\mathrm{i}} \epsilon^{\mathrm{i}}_{n;i,j}. \tag{10}$$

Specifically, we use the so-called catch model to characterise integrin detachment so that the maximum lifetime of the bound state occurs at intermediate values of $F^{\mathrm{i}}_{n;i,l}$. Larger forces lead to exponential decay in the lifetime of the bound state. This effect is captured by the following functional form of the integrin unbinding rate:

$$\mathrm{K}_{\mathrm{off}}(F^{\mathrm{i}}_{n;i,l}) = \lambda_1 \exp\Big(\gamma_1 F^{\mathrm{i}}_{n;i,l}\Big) + \lambda_2 \exp\Big(-\gamma_2 F^{\mathrm{i}}_{n;i,l}\Big). \tag{11}$$

Since integrin binding to ECM fibres occurs at a constant rate, $\mathrm{K}_{\mathrm{on}}$, the kinetics of the system is governed by the following system of ODEs:

$$\begin{aligned} \frac{dN^b_{n;i,l}}{dt} &= \mathrm{K}_{\mathrm{on}}\, N^u_{i,l} - \mathrm{K}_{\mathrm{off}}(F^{\mathrm{i}}_{n;i,l}) N^b_{n;i,l}, \\ \frac{dN^u_{i,l}}{dt} &= -\,\mathrm{K}_{\mathrm{on}}\, N^u_{i,l} + \mathrm{K}_{\mathrm{off}}(F^{\mathrm{i}}_{n;i,l}) N^b_{n;i,l}. \end{aligned} \tag{12}$$

Assuming conservation of the total number of available integrins in a FA site, we can reduce the binding-unbinding process (12) to one equation for the number of bound integrins, given by:

$$\frac{dN^b_{n;i,l}}{dt} = \mathrm{K}_{\mathrm{on}}\, N_{n;i,l} - \Big(\mathrm{K}_{\mathrm{on}} + \mathrm{K}_{\mathrm{off}}(F^{\mathrm{i}}_{i,l})\Big)\, N^b_{n;i,l}(t). \tag{13}$$

### S.3 Full system of model equations and parameter values

Our model consists of the three building blocks described above: a model describing the ECM fibre network, a cell model accounting for structural and active forces, and a cell-ECM interaction incorporated through the formation of FAs. We will now summarise key aspects of each modelling component and the coupling between them.

#### Model equations

The complete set of equations describing the evolution of the cell-ECM system is given as follows:

$$\begin{aligned}
&\eta \frac{d\mathbf{x}_l^{\mathrm{e}}}{dt} = \mathbf{F}_l^{\mathrm{el}} - \mathbf{F}_{l,n;i}^{\text{e-c}} \\
&\eta \frac{d\mathbf{x}_0^{\mathrm{c}n}}{dt} = \sum_{i \in \Omega_n^{\mathrm{c}}} -\mathbf{F}_{n;i}^{\mathrm{st}} - \mathbf{F}_{n;i}^{\mathrm{co}} \\
&\eta \frac{d\mathbf{x}_i^{\mathrm{c}n}}{dt} = \mathbf{F}_{n;i}^{\mathrm{st}} + \mathbf{F}_{n;i}^{\mathrm{co}} + \mathbf{F}_{n;i,l}^{\text{e-c}} \\
&\frac{dN_{n;i,l}^b}{dt} = \mathrm{K_{on}}\, N_{n;i,l} - \Big(\mathrm{K_{on}} + \mathrm{K_{off}}(F_{n;i,l}^{\mathrm{i}})\Big)\, N_{n;i,l}^b \\
&\text{for } l \in \Omega_{\mathrm{e}};\ \text{for } n = 1, \dots, N_{\mathrm{c}};\ \text{for } i \in \Omega_n^{\mathrm{c}}.
\end{aligned} \tag{14}$$

The forces and parameters appearing in these equations are summarised in Tables 2 and S1 Table, respectively.

**Table 2. Forces in the model.**

| | |
|---|---|
| **ECM forces** | |
| $\mathbf{F}_l^{\mathrm{el}}(t) = \sum_{m \in \Omega_{\mathrm{e},l}} -k_{l,m}(\lVert \mathbf{x}_l^{\mathrm{e}} - \mathbf{x}_m^{\mathrm{e}} \rVert - L_{l,m}^0) \frac{\mathbf{x}_l^{\mathrm{e}} - \mathbf{x}_m^{\mathrm{e}}}{\lVert \mathbf{x}_l^{\mathrm{e}} - \mathbf{x}_m^{\mathrm{e}} \rVert}$ | **Elastic force** of ECM fibre applied to crosslink $l$ |
| $\Omega_{\mathrm{e},l}$ | Set of crosslinks that are connected with $\mathbf{x}_l^{\mathrm{e}}$ |
| **Cell variables** | |
| $\mathbf{F}_{n;i}^{\mathrm{st}} = \mathbf{F}_i^{\mathrm{ve}} + \mathbf{F}_{n;i}^{\mathrm{an}}$ | **Structural forces** of the cell $n$ at the $i$-th adhesion site |
| $\mathbf{F}_i^{\mathrm{ve}} = (-\kappa_i^{\mathrm{el}} \varepsilon_i^{\mathrm{c}} - \kappa_i^{\mathrm{vi}} \dot{\varepsilon}_i^{\mathrm{c}}) \frac{\mathbf{x}_{n;i}^{\mathrm{c}} - \mathbf{x}_{n;0}^{\mathrm{c}}}{\lVert \mathbf{x}_{n;i}^{\mathrm{c}} - \mathbf{x}_{n;0}^{\mathrm{c}} \rVert}$ | Viscoelastic structural cellular force |
| $\varepsilon_i^{\mathrm{c}} = \lVert \mathbf{x}_{n;i}^{\mathrm{c}} - \mathbf{x}_{n;0}^{\mathrm{c}} \rVert - l_i^{\mathrm{c}}$ | Stretch of viscoelastic element $i$ |
| $\mathbf{F}_i^{\mathrm{an}} = -\nabla_i \left( \sum_{j \neq i, j \in \Omega_{\mathrm{Cell}}} \frac{1}{2} \kappa_{i,j}^{\mathrm{be}} (\cos\theta_{i,j} - \cos\theta_{i,j}^0) \right)$ | Angular structural cellular force acting on $i$-th element |
| $\cos\theta_{i,j} = \langle \hat{\mathbf{x}}_i, \hat{\mathbf{x}}_j \rangle$ | Cosine of the angle between cell viscoelastic elements $i$ and $j$ |
| $\mathbf{F}_i^{\mathrm{co}} = -F^{\mathrm{co}} \frac{\mathbf{x}_{n;i}^{\mathrm{c}} - \mathbf{x}_{n;0}^{\mathrm{c}}}{\lVert \mathbf{x}_{n;i}^{\mathrm{c}} - \mathbf{x}_{n;0}^{\mathrm{c}} \rVert}$ | **Active cell contraction force** |
| **Interaction forces: force at FAs** | |
| $\mathbf{F}_{n;i,l}^{\text{e-c}} = -N_{n;i,l}^b (k^{\mathrm{i}} \epsilon_{n;i,l}^{\mathrm{i}}) \frac{\mathbf{x}_{n;i}^{\mathrm{c}} - \mathbf{x}_l^{\mathrm{e}}}{\lVert \mathbf{x}_{n;i}^{\mathrm{c}} - \mathbf{x}_l^{\mathrm{e}} \rVert}$ | **Interaction force** between cell's element $i$ and ECM crosslink $l$ |
| $\epsilon_{n;i,l}^{\mathrm{i}}(t) = \left( \lVert \mathbf{x}_{n;i}^{\mathrm{c}} - \mathbf{x}_l^{\mathrm{e}} \rVert - L_0^{\mathrm{i}} \right)$ | Stretch of the integrins |
| $F_{i,l}^{\mathrm{i}} = k^{\mathrm{i}} \epsilon_{i,j}^{\mathrm{i}}$ | Total force exerted on an integrin at FA connecting cell element $i$ and ECM crosslink $l$ |

#### Initial conditions

In order to solve the system of equations given by Eq (14), we have to determine an initial configuration of the system. At first, we generate an arbitrary isotropic ECM configuration, with no preferential alignment or stresses at the fibres. The details for the ECM generation are given in Section S.4. The cells are positioned on the ECM so that their adhesion sites $\mathbf{x}^{\mathrm{c}}_{n;i}$, $i = 1, \dots, N^{\mathrm{c}}_n$ are attached to their closest ECM crosslinks. The number of bound integrins attached and their stretch at the different FAs are adjusted to be close to equilibrium when no active forces act on the system.

### S.4 Numerical methods

In this section we summarise the numerical methods utilised in this work.

#### Initial condition generation:

1. In order to generate different random initial ECM configurations, we employ the method proposed in [1]. Given the density of ECM fibres, we randomly distribute the corresponding number of seed points within the 2D domain. We refer to [1] for the description of the number of seed points. The ECM network is then obtained by generating a Voronoi tessellation on the set of seed points. We assume that the vertices of the Voronoi tesselation correspond to the ECM crosslinks and its edges represent ECM fibres.
2. Once the ECM configuration is generated, we place on it cells with the desired shape. We then determine the closest crosslink to the cell adhesion sites which defines the cell-ECM attachment.
3. The cells are deformed to ensure that the distance between the cell adhesion site and closest crosslink is on the order of the natural length of an integrin. We initialise each FA with a certain number of bound integrins to facilitate FA formation. In this work, we start with 10 bound integrins.
4. Lastly, we let the system relax. In the relaxation process we consider no active contraction forces. The resulting equilibrium configuration corresponds to the initial condition we use in our model simulation. At the initial configuration we have a certain number of bound integrins at each FA.

**ODE approximation:** For the numerical solution of the system of equations given by Eq (14), we use the Runge-Kutta solver `ode45` implemented in Matlab. Since the system is stiff, we use a time step of the order of $10^{-8}$. For the system given by Eq (9) in main text, we use the implicit Matlab solver `ode15s` for stiff systems.

**Bifurcation diagram:** To obtain the bifurcation diagrams shown in Fig 5, we use the Bifurcation Kit package from Julia [18].

### S.5 Exact calculation of the critical value $F^{\mathrm{BP}}$

Fig 5 shows that the system exhibits a saddle-node bifurcation. At this point, the nullclines (curves defined below) intersect at a single point, $(\epsilon_b^{\mathrm{BP}}, N_b^{\mathrm{BP}})$, and they are tangent to each other. In this section, we describe how to calculate exactly the critical value of the external force, $F^{\mathrm{BP}}$, corresponding to the saddle-node bifurcation in Eq (9) in main text. We obtain an implicit expression for the critical force, the proportion of the bound integrins $N_b^{\mathrm{BP}}/N$ and their stretch $\epsilon_b^{\mathrm{BP}}$ which can be solved numerically. The detailed step-by-step derivation is described below.

1. We define the nullclines of the system given by Eq (9) in main text, as pairs $(\epsilon_b, N_b)$ such that either $\frac{d\epsilon_b}{dt} = 0$ or $\frac{dN_b}{dt} = 0$. The points of intersection of these two nullclines can be calculated by solving the following equation:

$$\frac{N_b}{N} = \frac{\mathrm{K_{on}}}{\mathrm{K_{on}} + \mathrm{K_{off}}(k^{\mathrm{i}}\epsilon_b)} = \frac{\mathcal{F}}{N}\frac{1}{2k^{\mathrm{i}}\epsilon_b} \tag{15}$$

2. Furthermore, the condition for these nullclines to be tangent to each other requires that the following expression is satisfied:

$$-\frac{k^{\mathrm{i}}\,\mathrm{K_{off}}'(k^{\mathrm{i}}\epsilon_b)}{(\mathrm{K_{on}} + \mathrm{K_{off}}(k^{\mathrm{i}}\epsilon_b))^2} = -\frac{\mathcal{F}}{N}\frac{2k^{\mathrm{i}}}{(2k^{\mathrm{i}}\epsilon_b)^2}, \tag{16}$$

where $\mathrm{K_{off}}'(k^{\mathrm{i}}\epsilon_b)$ refers to the derivative of the ratio of unbound evaluated at the force exerted on each integrin.

3. Using Eqs (15)-(16), we obtain an equation for the steady-state value of the stretch at the critical point:

$$\epsilon_b^{\mathrm{BP}} = \frac{\mathrm{K_{on}} + \mathrm{K_{off}}(k^{\mathrm{i}}\epsilon_b^{\mathrm{BP}})}{k^{\mathrm{i}}\,\mathrm{K_{off}}'(k^{\mathrm{i}}\epsilon_b^{\mathrm{BP}})}. \tag{17}$$

The solution of Eq (17) can be calculated numerically. We use the `fzero` Matlab function to obtain $\epsilon_b^{\mathrm{BP}}$.

4. From Eq (15), and $\epsilon_b^{\mathrm{BP}}$, we can calculate the critical force value, $F^{\mathrm{BP}}$, beyond which no stable equilibria exist:

$$\frac{F^{\mathrm{BP}}}{N} = \frac{2k^{\mathrm{i}}\epsilon_b^{\mathrm{BP}}\,\mathrm{K_{on}}}{\mathrm{K_{on}} + \mathrm{K_{off}}(k^{\mathrm{i}}\epsilon_b^{\mathrm{BP}})}. \tag{18}$$

5. Lastly, we determine the proportion of bound integrins at the equilibrium at the bifurcation point:

$$\frac{N_b^{\mathrm{BP}}}{N} = \frac{\mathrm{K_{on}}}{\mathrm{K_{on}} + \mathrm{K_{off}}(\epsilon_b^{\mathrm{BP}})}. \tag{19}$$

From Eqs (17) and (19), we notice that the value of the equilibrium stretch and the proportion of bound integrins at the bifurcation point are independent of the value of the total available integrins in a focal adhesion, $N$. This is illustrated in Fig 5B.
