## Supplementary material for "Multiscale modelling shows how cell-ECM interactions impact ECM fibre alignment and cell detachment": S1 Table

**Table 1. Model parameters.** Description and values of the parameters used in our simulations.

| | |
|---|---|
| **General parameters** | |
| $\eta = 50$ pN·s·$\mu$m$^{-1}$ | Viscous coefficient. Estimated |
| **ECM parameters** | |
| $L^0_{l,m}$ | Natural length of the fibre.<br>Determined by the initial condition |
| $k^{\mathrm{el}}_{l,m} = \frac{E_{l,m} A_{l,m}}{L^0_{l,m}}$ | Elastic constant of the fibre |
| $E_{l,m}$<br>$E_{l,m} = E_{\mathrm{t}} = 10 - 100$ MPa<br>$E_{l,m} = E_{\mathrm{c}} = 0.1 E_{\mathrm{t}}$ MPa | Young's modulus of the fibres. Reference value from [1,2]<br>Young's modulus of extended fibres<br>Young's modulus of compressed fibres [2] |
| $A_{l,m}$<br>Fibre diameter $= 10^{-7}$ m | Cross-sectional area of the fibres<br>[3,4] |
| Crosslink density, $0.2\,\mu\mathrm{m}^{-2}$ | [2] |
| Mean fibre density, $0.6 - 0.7\,\mu\mathrm{m} \cdot \mu\mathrm{m}^{-2}$ | [2] |
| **Cell parameters** | |
| $\kappa^{\mathrm{el}}_i = 5 \cdot 10^3$ pN·$\mu$m$^{-1}$ | Elastic constant of cell viscoelastic elements. Estimated |
| $\kappa^{\mathrm{vi}}_i = 10^3$ pN·s·$\mu$m$^{-1}$ | Damping constant of cell viscoelastic elements. Estimated |
| $l^{\mathrm{c}}_i = 15$ $\mu$m | Natural length of viscoelastic elements.<br>Estimated from cell sizes. |
| $\kappa^{\mathrm{el}}_i = 1$ pN·rad$^{-1}$ | Angular elastic constant of the viscoelastic elements.<br>Estimated |
| $\theta^0_{i,j}$ | Natural angle between viscoelastic elements.<br>Determined by initial cell shape. |
| $F^{\mathrm{co}} = 10^3 - 5 \times 10^4$ pN | Estimated. Reference values [5–7]. |
| **Integrin and FAs parameters** | |
| $k^{\mathrm{i}} = 10^6$pN $\cdot \mu$m$-1$ | Elastic constant of the integrins, see [8]. |
| $\mathrm{K_{on}} = 0.2\,\mathrm{s}^{-1}$ | Integrins binding rate, estimated value.<br>Reference value for estimation [7,9] |
| $\mathrm{K_{off}}(F^{\mathrm{i}}_{i,l}) = 0.4 \exp\left(-0.04 F^{\mathrm{i}}_{i,l}\right) + 4 \times 10^{-7} \exp\left(0.2 F^{\mathrm{i}}_{i,l}\right)$ | Integrin unbinding rate, [9] |
| $N = \frac{6000}{N^{\mathrm{c}}_n}$ integrins | Estimated available integrins per FA site.<br>$N^{\mathrm{c}}_n$ is the number of FA sites in the $n$-th cell |
